# How AlphaFold and related models predict protein-peptide complex structures

**DOI:** 10.1101/2025.06.18.660495

**Authors:** Lindsey Guan, Amy E. Keating

## Abstract

Protein-peptide interactions mediate many biological processes, and access to accurate structural models, through experimental determination or reliable computational prediction, is essential for understanding protein function and designing novel protein-protein interactions. AlphaFold2-Multimer (AF2-Multimer), AlphaFold3 (AF3), and related models such as Boltz-1 and Chai-1 are state-of-the-art protein structure predictors that successfully predict protein-peptide complex structures. Using a dataset of experimentally resolved protein-peptide structures, we analyzed the performance of these four structure prediction models to understand how they work. We found evidence of bias for previously seen structures, suggesting that models may struggle to generalize to novel target proteins or binding sites. We probed how models use the protein and peptide multiple sequence alignments (MSAs), which are often shallow or of poor quality for peptide sequences. We found weak evidence that models use coevolutionary information from paired MSAs and found that both the target and peptide unpaired MSAs contribute to performance. Our work highlights the promise of deep learning for peptide docking and the importance of diverse representation of interface geometries in the training data for optimal prediction performance.

## Introduction

Many protein-protein interactions involve a well-folded domain engaging a smaller peptide [1]. However, the Protein Data Bank (PDB) captures only a small fraction of these interactions, which are often low-affinity and dynamic. Protein structure-prediction methods, such as AlphaFold2 (AF2), AF2-Multimer, and AlphaFold3 (AF3) [2, 3, 4], can accurately predict many protein-protein complex structures, and benchmarks for protein-peptide docking have shown that performance is also promising for these complexes. AF2, which was trained on only single-chain structures, predicted 51% of domain-motif complexes with < 2.5 Å interface backbone RMSD when “hacked” to predict the structure of complexes by fusing a protein to its peptide partner using a poly-glycine linker [5]. Ko et al. found that 50% of protein-peptide complexes, fused with a glycine linker, were predicted with < 2 Å peptide backbone RMSD [6]. AF2-Multimer, which was trained on single chains and multi-chain complexes, encodes chains separately and predicts structures for complexes without requiring a linker. Johansson-Åkhe et al. found that AF2-Multimer outperformed other established methods and achieved an acceptable prediction in 60% of test cases [7]. A summary of methods and success rates reported in previous studies can be found in Table S1.

Because peptides are shorter than globular proteins and evolve under functional constraints that can be difficult to capture in multiple-sequence alignments (MSAs) [8, 9], they represent a special case for models that heavily rely on an MSA. Since the release of AF2, structure prediction models have been developed that do not use an MSA, such as ESMFold and OmegaFold [10, 11]. However, MSA-based models remain the most performant for predicting protein complex structures [12]. Bryant et al. [13] observed that AF2-Multimer predicted bacterial protein-protein complexes more accurately than complexes from eukaryotes, which they attributed to having more sequenced bacterial genomes to search from [14]. They observed that the fraction of correctly predicted structures increased with a larger number of non-redundant sequences in the MSA. For predictions involving multiple chains, cross-chain coevolutionary information is available to structure prediction models through sequence pairing during MSA construction. Sequences from the same species are paired together in each row of the MSA, followed by the unpaired sequences on the block diagonal. Whether and how AF2-Multimer and newer models use the MSA input, including MSA pairing, to make accurate predictions for protein-peptide complexes remains unclear.

Structure prediction models form the basis of many methods developed for the discovery or design of protein-binding peptides [15, 16, 17, 18]. To inform further development of such methods, we tested the requirements for accurate deep learning-based protein-peptide complex predictions. We benchmarked AF2 with monomer weights (AF2), AF2-Multimer, AF3, Boltz-1, and Chai-1 on a set of protein-peptide complexes, reproducing and extending earlier findings of good performance [3, 4, 19, 20]. We then sought to explain the observed range of prediction accuracies through an assessment of sequence and structural similarity to the training set and found a strong dependence of prediction accuracy on target-training overlap. We investigated the ability of models to process MSA statistics and found that inter-chain coevolution is weakly related to prediction success for protein-peptide and protein-protein complexes. However, we also showed that the information in the unpaired MSAs for both protein and peptide chains contributes to prediction accuracy. Strikingly, good predictions can be made for many complexes when the peptide sequence is masked. The target MSA contains information that guides binding-site prediction, and the peptide MSA improves both binding-site and pose prediction.

## Results

### State-of-the-art methods accurately predict most protein-peptide complex structures

To curate a set of protein-peptide complexes, we filtered structures from the PDB by peptide length, presence of cofactors, and presence of crystal contacts near the peptide binding site (Figure 1a) [21]. We also clustered the target and peptide sequences at 95% sequence identity. We made predictions with unpaired MSAs and no templates, took the top-ranked model out of five predictions, and calculated DockQ values between predicted and native structures. DockQ is a linear combination of the peptide backbone RMSD, interface backbone RMSD, and number of recovered native contacts normalized to the range 0-1, with 1 representing perfect recovery [22, 23]. In describing docking accuracy, we refer to the performance categories described by Basu et al., which map closely to the “acceptable”, “medium”, and “high-quality” predictions used in the CAPRI assessment [24]. For all methods, many are high quality (DockQ > 0.80), with the peptide binding pose mostly correct (Figure 1b, Figure S1a). This performance is consistent with previous studies benchmarking AF2-Multimer for protein-peptide structure prediction (Table S1) [5, 6]. Some predictions are medium (DockQ > 0.49) or poor (DockQ < 0.23); in these cases, the peptide orientation or the binding site is incorrect.

**Figure 1.**
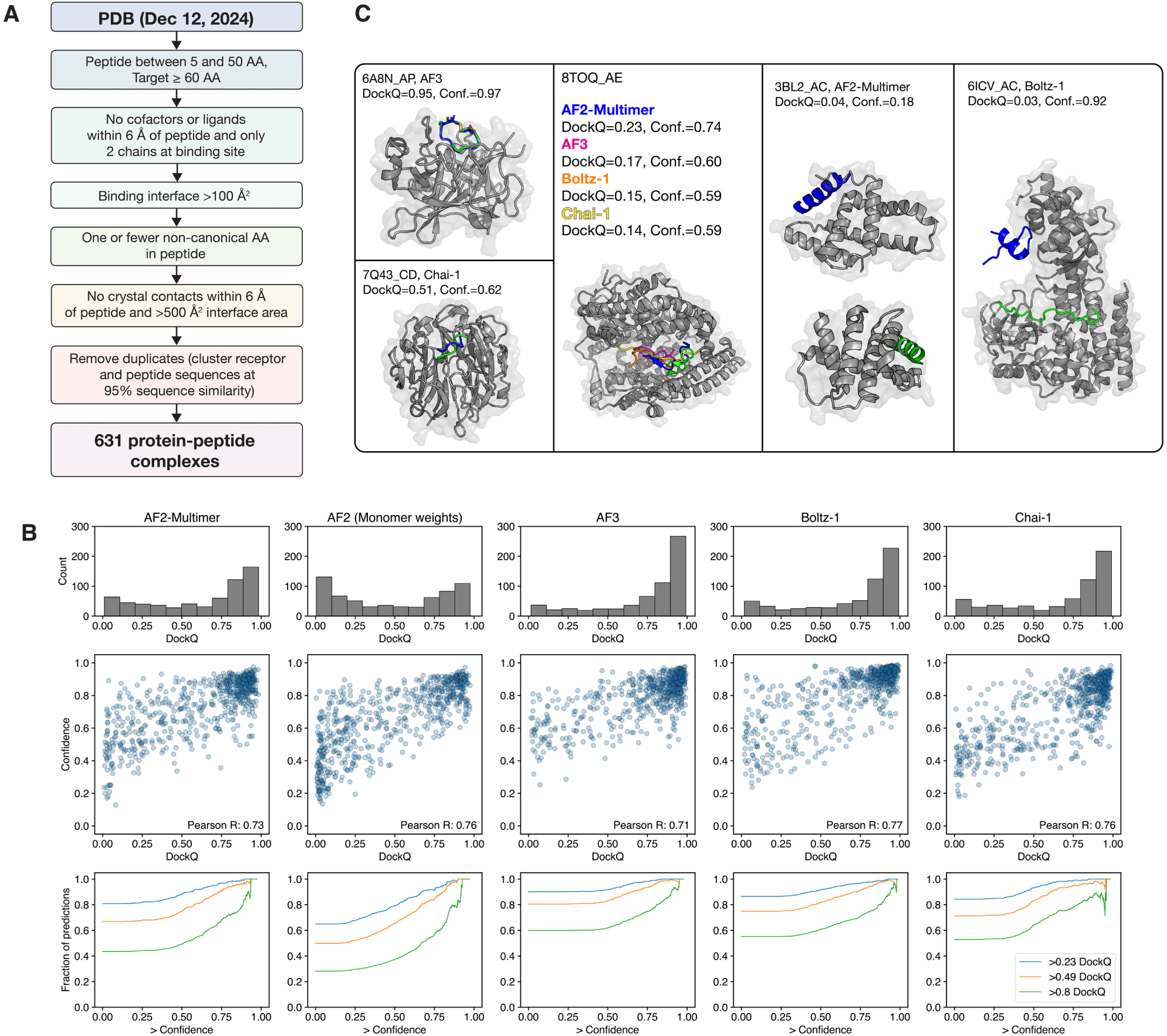
Structure prediction methods accurately dock most test-set peptides. **(a)** Curating protein-peptide complexes. Applying filters and clustering resulted in a test set of 631 protein-peptide complexes. **(b)** top: Model confidence and accuracy of all models on the test-set structures as measured by confidence (ipTM+pTM) and DockQ; bottom: Cumulative distributions showing DockQ as a function of ipTM+pTM. The vertical axis tracks the fraction of samples with DockQ greater than the value indicated in the legend, as a function of the confidence cutoff indicated on the horizontal axis. **(c)** Example predictions. For 6A8N_AP, 7Q43_CD, 3BL2_AC, and 6ICV_AC, predictions are in blue and native peptides are in green. For 8TOQ_AE, predictions are in blue, magenta, orange, and yellow, and the native peptide is in green. For all except 3BL2_AC, only the native target structure is shown.

Five complexes predicted with varying accuracy are shown in Figure 1c. PDB 6A8N chains A and P (henceforth 6A8N _AP) provide an example of a highly accurate prediction with AF3: the model captured the correct binding interaction and the disulfide bond cyclizing the peptide [25, 26]. The structure predicted for 7Q43_CD using Chai-1 is a medium accuracy prediction where the binding site is correct, but the peptide orientation is not [27]. For 8TOQ_AE, all models predicted the peptide orientation incorrectly, leading to low DockQ [28, 29]. There are a few instances of models failing to capture the target structure, such as the prediction of 3BL2_AC with AF2-Multimer [30, 31]. Finally, many instances of poor predictions are the result of an incorrect binding-site prediction, such as the high-confidence yet incorrect prediction of structure 6ICV_AC by Boltz-1 [32, 33]. In addition to DockQ, which uses backbone-only RMSD calculation, we also defined criteria to reflect what we refer to as atomically accurate predictions: > 90% of native contacts recovered, no clashes, peptide all-atom RMSD < 2 Å, and interface all-atom RMSD < 3 Å. 16-40% of predictions were accurate to this degree, with AF3 achieving the best performance (Figure S1b).

The AF2-Multimer confidence score is reported to be an excellent scoring function for multimeric structure predictions. Models that used the highest-ranking prediction after extensive structural sampling surpassed other methods at CASP15 [34]. For our protein-peptide test set, the Pearson correlation coefficient between confidence (ipTM+pTM) and DockQ was > 0.73 for all methods. At ipTM+pTM > 0.75, 70% of predictions are high-quality and 90% are medium- or high-quality across all models (Figure 1b, bottom). We also calculated pDockQ [13], which yielded a worse correlation with DockQ than ipTM+pTM (Figure S2).

To understand how well models predict the secondary structure adopted by the bound peptide, irrespective of docking accuracy, we used DSSP to define secondary structure in the native and predicted peptide structures [35, 36]. All models predicted more residues to be *α*-helical and less loop-like than the experimental structures (Figure S3a). Complexes where the native interface is composed mostly of loop-like residues were, on average, predicted less successfully than complexes with regular secondary structure (Figure S3b).

### Prediction accuracy is lower for previously unseen complexes

We tested how prediction accuracy relates to the similarity of the target complex to the training set. Previous work reported similar AF2 performance for protein-peptide structures released before and after the AF2 training cutoff [5, 18]. To examine whether there is a dependency for AF3, Boltz-1, and Chai-1, we constructed a post-training cutoff set and a pre-training cutoff set, each consisting of 74 structures, as described in the methods (Figure 2a). The difference in prediction accuracy between the test-set and training-set structures was significant for all models (*p* = 0.025 to 6.4E-4; Figure 2b), except AF2 with monomer weights (*p* = 0.6), which is expected because this training set did not include any complexes. When evaluating the models on only post-cutoff structures, all models, except AF2 with monomer weights, had comparable performance (with median DockQ ranging from 0.64 to 0.69). Strikingly, with AF3, Boltz-1 and Chai-1, only 7-16% of post-cutoff predictions were atomically accurate, while 23-39% of pre-cutoff predictions were atomically accurate (Figure S4b).

**Figure 2.**
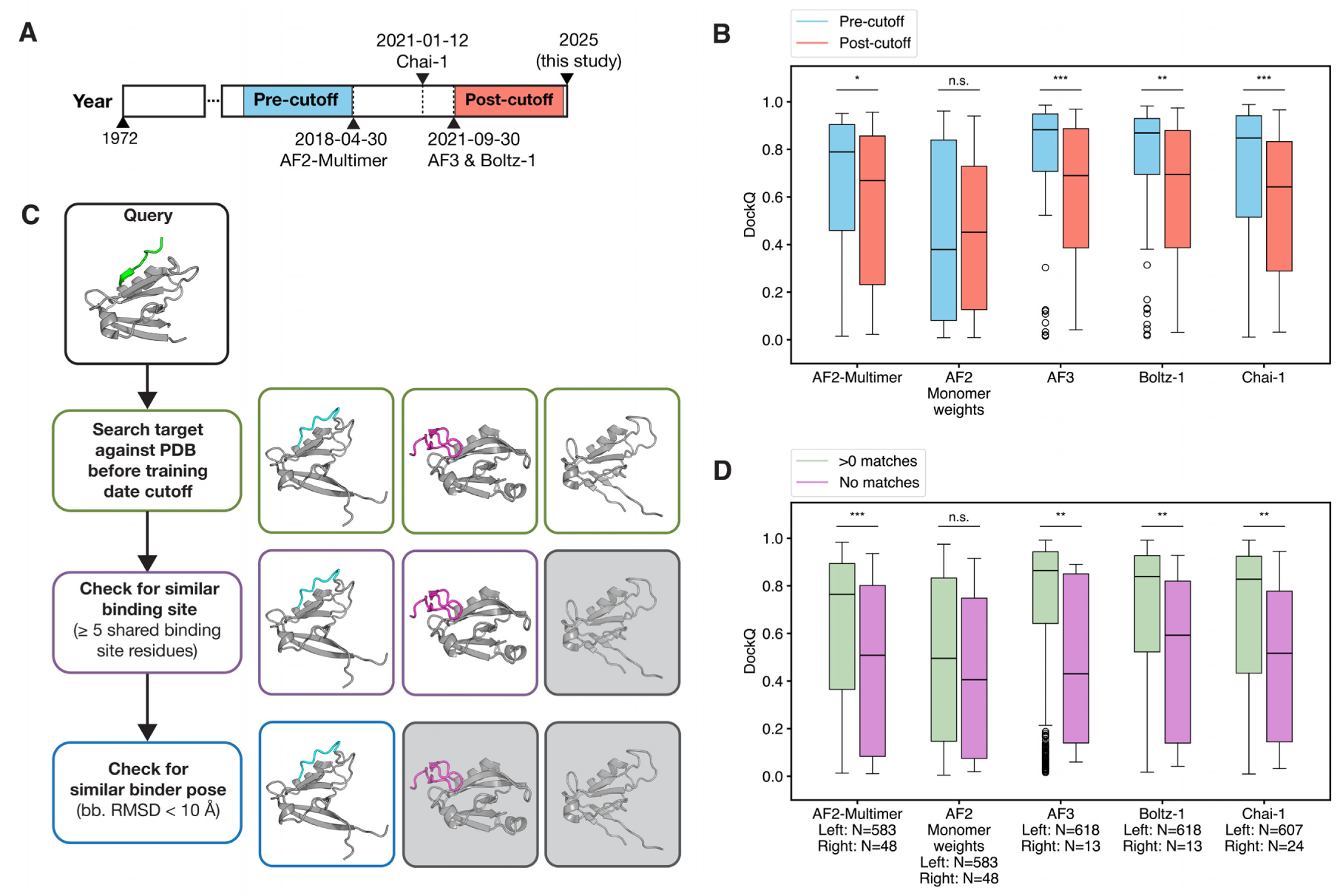
Prediction accuracy improves with similarity to training set examples. **(a)** Sets of pre-cutoff and post-cutoff structures were selected based on the training dates for the different models and the PDB structure release date. **(b)** Distribution of DockQ scores for predictions where the PDB entry was seen during training (pre-cutoff) and where it was unseen (post-cutoff), N=74 for each. **(c)** Procedure for identifying binding-site matches for a given complex in the PDB (details in Methods). Structures on the right are hits at each step in the pipeline, with greyed-out structures filtered out. **(d)** Distribution of DockQ for predictions where the binding interaction was represented in the training set vs. not represented in the training set. (two-sided Mann-Whitney U-test, \**p* < 0.05, \*\**p* < 0.01, \*\*\**p* < 0.001).

In a more stringent assessment of similarity between the training and test examples, for each test complex, we searched for homologous targets (by sequence or structure) in the training set and checked whether matching structures were bound to a partner structurally similar to the peptide (< 10 Å RMSD) (Figure 2c, Methods). If so, we designated the training set complex as a binding-site match to the target complex. For all models, binding sites that were represented by at least one complex in the training set were predicted more accurately than binding interactions that were not represented (two-sided Mann-Whitney U-test, *p* < 0.05, Figure 2d), with the difference being least significant for AF2 with monomer weights. Although AF2 with monomer weights did not exhibit the same level of bias to the training set as the other models, performance was worse in most splits, and we did not include this method in subsequent analyses.

We saw little correlation between the number of binding-site matches in the training set and DockQ, although highly represented complexes were almost always well predicted. Across models, 92-100% of complexes with more than 200 binding-site matches had DockQ *≥* 0.49 (Figure S4a). Among structures with no homologous interactions in the training set, none were atomically accurate with the AF3-like models (Figure S4b). This training-set bias does not arise from memorization of the bound peptide structure, since unbound peptide predictions tended to be dissimilar from the native bound peptide structure, with median TM scores of 0.28 - 0.38 (Figure S4c).

We also tested whether incorrect predictions corresponded to binding modes seen frequently in the training data. For a substantial fraction of cases (e.g., 46% for AF3), the predicted interface had more homologous matches in the training set than the native interface (Figure 3a). One example is the prediction of 8V8E_AC with AF2-Multimer and Boltz-1 [37, 38]. The predicted complex had 3 and 10 similar structures in the AF2-Multimer and Boltz-1 training sets, respectively, while the native complex had none (Figure 3b). The peptide was predicted to bind in a pocket represented more frequently as a peptide binding site. In 8V8E_AC, this binding pocket is occupied by a ligand, which raised the question of whether Boltz-1 would choose the correct site for the peptide if provided the ligand. However, when we provided Boltz-1 with both the peptide and ligand, it docked the ligand in the correct site and the peptide in another incorrect binding site (Figure 3b, right). In some cases, the predicted and native binding sites had a similar number of matches in the training set. For 6TYX_AC, the native and predicted binding sites had 4 and 6 matches, respectively [32, 39]. Among the 5 predictions from AF2-Multimer, the top-ranked and third-ranked predictions placed the peptide in the same incorrect binding site, while the second-, fourth-, and fifth-ranked structures placed the peptide in the native site (Figure 3c). We also found interesting instances where the target protein has at least two peptide-binding regions. For example, the ARM domain of importin *α* has two nuclear-localization-signal (NLS) binding sites, a major and minor site (Figure 3d) [40]. Both binding sites on 4B8P_A are well-represented in the training data, with 107 matches for the major site and 73 matches for the minor site. In 4B8P_AC, the NLS peptide binds specifically to the minor site of *Oryza sativa* importin *α*, though AF3 predicted the peptide to bind in the major site and, interestingly, docked the N-terminus of the ARM domain in the minor site (Figure 3d) [41, 42].

**Figure 3.**
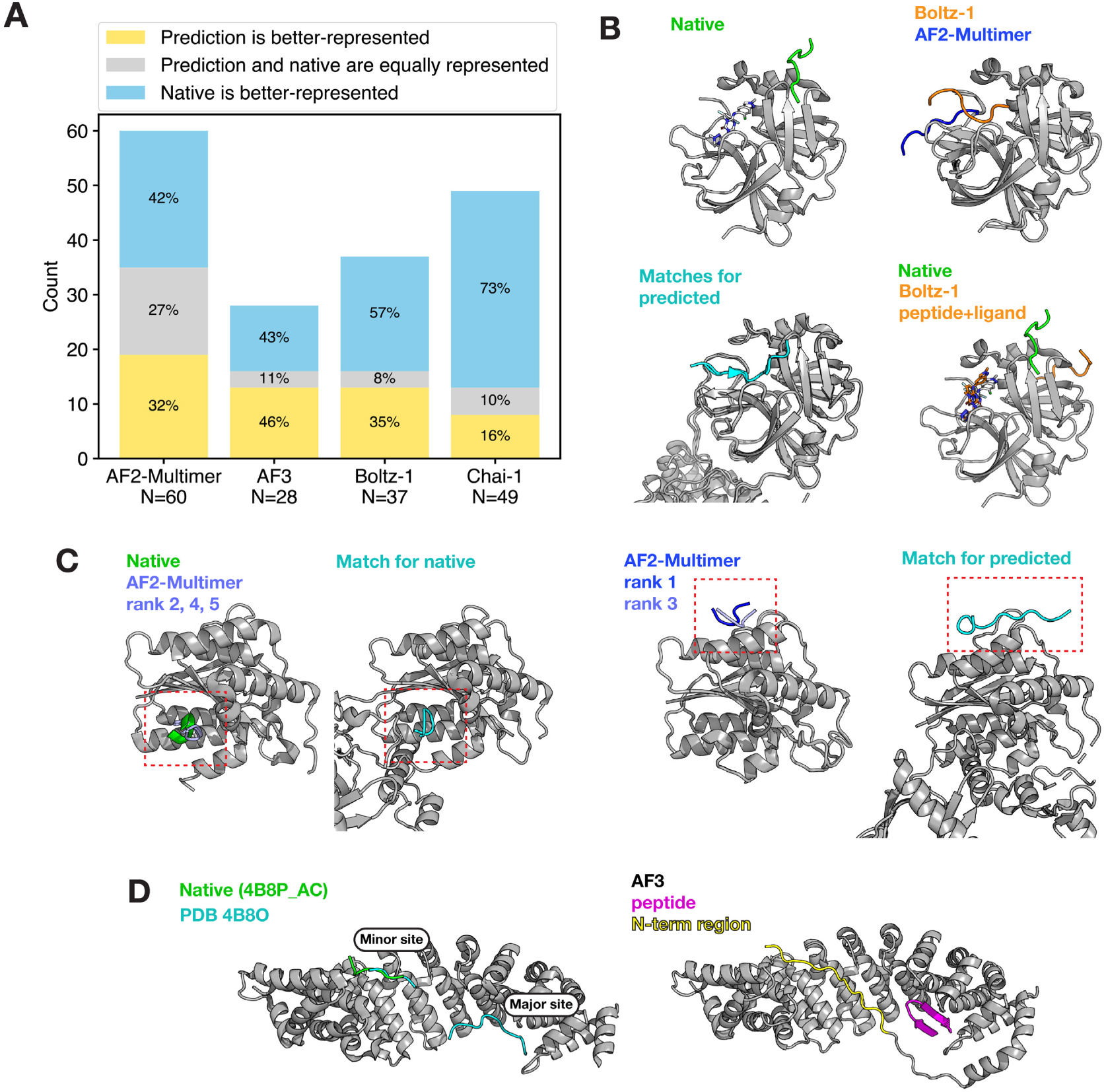
Some incorrect predictions may be explained by structures seen frequently in the training data. **(a)** Binding site representation for the native and predicted binding sites for poor predictions where DockQ < 0.23 and target TMscore > 0.6. **(b)** top left: native structure for 8V8E_AC; top right: predictions for the 8V8E_AC complex with Boltz-1 (conf. = 0.80) and AF2-Multimer (conf. = 0.72); bottom left: a representative subsample of binding site matches in the training data for the predicted structures, PDB 2Q6G, 3D23; bottom right: native and predicted structure when the ligand, ensitrelvir, is provided to Boltz-1 (conf.= 0.41). **(c)** left: native structure for 6TYX_AC with the second-, fourth-, and fifth-ranked AF2-Multimer predictions (conf. = 0.47, 0.44, 0.43, respectively); middle left: a representative binding site match for the native complex, PDB 6ERG; middle right: the top- and third-ranked AF2-Multimer predictions (conf. = 0.48, 0.46, respectively); right: a representative binding site match for the predicted complex, PDB 6ERF. **(d)** left: native structure for 4B8P_AC in green, with PDB 4B8O shown in cyan to illustrate the major and minor NLS-binding regions of importin *α*; right: AF3 prediction for 4B8P_AC.

### Models do not rely on MSA sequence pairing for protein-peptide predictions

Despite the strong association between high prediction accuracy and target-training set overlap, there were nevertheless correct predictions for complexes that lacked any binding-site matches in the training dataset (Figure 2d). There were also incorrect predictions for complexes with some homologous structures in the training data. This motivated the question of what information powers predictions across the entire test set. Previous studies have shown that MSA-based models rely on the MSA to make accurate predictions [2]. In our tests, too, models rarely made accurate protein-peptide structure predictions when we ablated the MSA, because the target could not be predicted accurately (Figure S5a). Most structures for which accurate predictions were made without an MSA were represented in the training set. The few novel complexes that were predicted correctly had interfaces with canonical secondary structure elements (e.g., the peptide forms a strand in a *β*-sheet). One was a *de novo* designed complex, 8FG6_BA [43], consistent with previous reports that *de novo* designed proteins can be predicted without an MSA (Figure S5b) [44, 45, 46].

We explored the importance of the peptide MSA and MSA pairing by species for structure prediction. Although most complexes had a deep MSA for the target, most peptide sequences had sparse peptide MSAs. 213/631 complexes had a non-empty peptide MSA, and 146/631 had a peptide MSA with at least 50 sequences (Figure S6a). The results described so far used these default MSAs without pairing by species and thus without access to protein-peptide coevolutionary information. When we tried to construct paired MSAs using default methods, only two complexes resulted in a paired MSA with at least 50 sequences; for most complexes, the paired MSA was empty (Figure S6a).

To test whether models can use coevolutionary information, we constructed deeper paired MSAs for cases where the peptide could be mapped to a canonical UniProt entry. Using SIFTS annotations, we extracted the protein sequence from which the peptide sequence was derived and included 50 or 100 flanking residues (“context”) around the PDB-delimited peptide [47]. These extended sequences were used as the queries when building paired MSAs. Then, for the peptide chain, all MSA columns were removed except those corresponding to the PDB-delimited peptide before the paired MSA was used as input to the models (Figure 4a). This procedure is similar to approaches used for other protein fragment binding predictions and yielded deeper paired MSAs for many complexes (Figure S6a) [17, 18, 48]. Shallow or low-quality MSAs constructed in this manner, as measured by Jensen-Shannon divergence, were filtered out (see Methods) [49], leaving 92 populated paired MSAs with 50 residues of context and 160 populated paired MSAs with 100 residues of context. To reduce variability in structure predictions caused by different unpaired MSAs samples across recycles, we only used the paired MSA.

**Figure 4.**
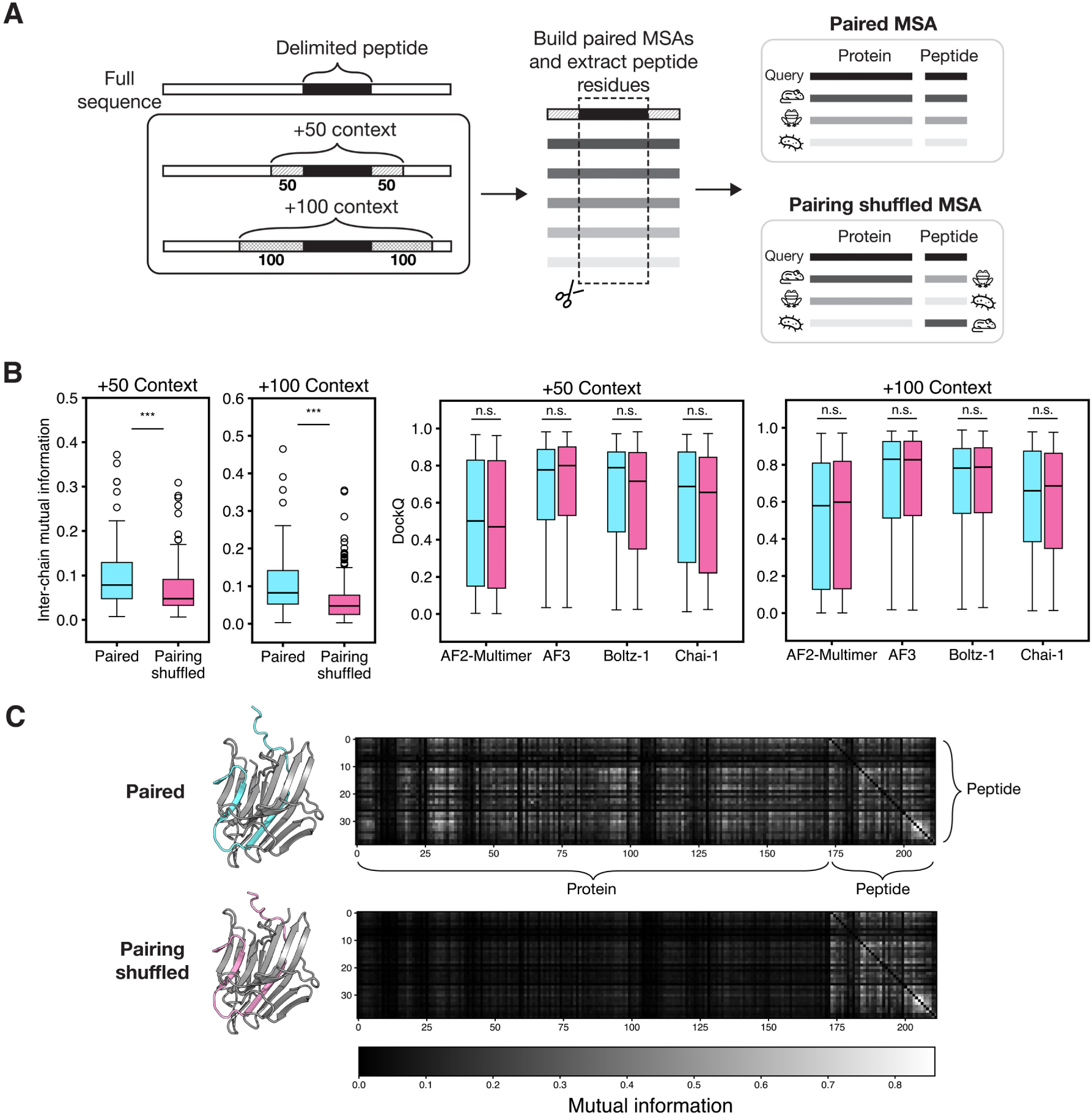
MSA pairing does not improve prediction performance. **(a)** Procedure for constructing MSAs using additional peptide context. Paired and shuffled MSAs were constructed using the peptide sequence with an additional 50 or 100 residues on either side of the delimited peptide. **(b)** Distribution of MI and DockQ scores for predictions made with paired (cyan) vs. shuffled paired (pink) MSAs, N = 92 for 50 residues of context and N = 160 for 100 residues of context (Wilcoxon signed-rank test, \**p* < 0.05, \*\**p* < 0.01, \*\*\**p* < 0.001). **(c)** An example is shown for 3TRS_BA with 100-residue context, predicted with AF2-Multimer. The structure prediction is shown for the paired MSA (cyan, DockQ = 0.86) and shuffled MSA (pink, DockQ = 0.82). MI for peptide residues with respect to every other residue in the query (target + peptide) is shown in the heatmap.

We compared prediction performance using the paired MSAs to predictions made using MSAs containing the same sequences but with protein-peptide pairings randomized. We calculated inter-chain mutual information (MI) for the original and shuffled alignments [50, 51, 52]. As expected, the shuffling procedure lowered the MI in the paired MSAs (Figure 4b, left). However, there was little difference in prediction accuracy between models, indicating that the coevolutionary signal between chains is not important for successful docking. In Figure 4c, for complex 3TRS_BA, the heavy and light chain of a peptidase from the fungus *Aspergillus niger* illustrates these trends: both the paired and shuffled MSA predictions identified the correct binding site, even when the MI between the peptide and target MSA columns decreased (Figure 4c, Figure S6) [53, 54]. Predictions made with MSA pairings based on 100 residues of context tended to be more accurate than predictions made with MSA pairings based on 50 residues of context, likely due to the deeper MSAs (Figure S6a).

To compare this result using deeper paired MSAs for protein-peptide complexes with data for protein-protein complexes, which typically have deeper and higher quality MSAs, we repeated the MSA shuffling procedure for a set of protein-protein complexes from Bryant et al. [13]. We focused on 340 complexes that yielded paired MSAs containing at least 50 sequences. Both the paired and unpaired MSAs were included in the prediction to ensure that the individual chains folded well (Figure S8b). MSA shuffling resulted in lower inter-chain mutual information (Figure S8a) and yielded slightly worse performance for AF2-Multimer and Boltz-1, while little effect was observed with AF3 or Chai-1 (Figure S8b). Although the decreases in performance for AF2-Multimer and Boltz-1 were statistically significant, our results show that ablating coevolution is only detrimental for a small subset of complexes (Figure S8c). One example of a complex for which the shuffling degraded performance is 4ZHY_BA, for which AF2-Multimer failed to recover the correct docked pose after the paired MSA was shuffled (Figure S8d) [55, 56].

### The roles of the unpaired target and peptide MSAs

One hypothesis for how deep-learning models dock peptides onto proteins is that models have learned to identify binding sites on the target. AF2 has previously been adapted for predicting small-molecule ligand binding sites on target protein [57]. We tested whether structure-prediction models have learned to predict binding sites irrespective of the peptide sequence that is provided. First, we tried ablating the peptide sequence using “unknown” tokens, “X” (Figure 5a). Chai-1 and Boltz-1 output a backbone structure for masked residues, which we compared to the native structure using a scaled peptide RMSD (Figure 5b). A lower RMSD_map_ value corresponds to a better prediction. We considered a scaled peptide RMSD score of < 2 Å an acceptable prediction (Figure S9). AF2-Multimer does not return a structure for a peptide composed of unknown tokens, so we analyzed and compared contact maps predicted by the distogram head using a new metric, “RMSD_map_” (Figure 5a). We considered RMSD_map_ < 3 a docking success because at this cutoff, 92% of unmasked peptides were predicted with DockQ > 0.23 (Figure S9).

**Figure 5.**
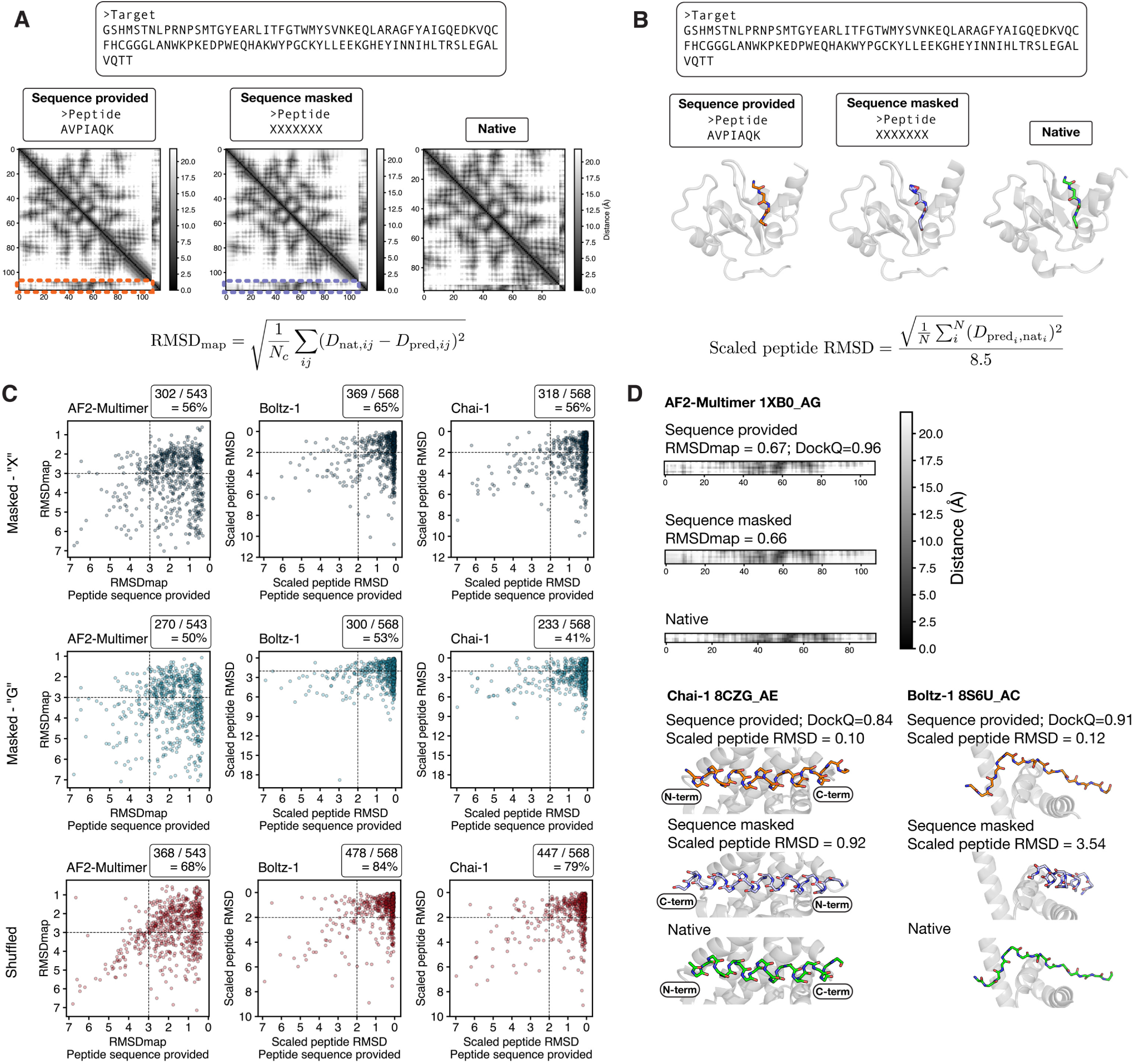
Peptides can be docked without knowledge of the peptide sequences. **(a)** Procedure for masking the peptide sequence and calculating the distance between native and predicted contact maps (RMSDmap) for AF2-Multimer predictions. In the contact maps, interchain contacts are shown in dashed boxes. 1XB0_AG is shown as an example [83, 84]. **(b)** Procedure for masking the peptide sequence and calculating scaled peptide RMSD for Boltz-1 and Chai-1. RMSD is calculated on peptide backbone residues after aligning the target. **(c)** RMSDmap and scaled peptide RMSD values for predictions made with the peptide sequence provided (x-axis) and with the peptide sequence modified in three ways (y-axes): masked with the token “X,” assigned as poly-glycine, or randomly shuffled. Fractions reflect the number of successful predictions that stayed successful after each sequence modification (i.e., examples in the upper-right section over those exceeding the indicated horizontal axis cutoff) **(d)** top: An example of the RMSDmap score and the corresponding interchain contact map; bottom: Two examples of the scaled peptide RMSD score and the corresponding prediction with peptide backbone residues are shown [85, 86, 87, 66].

AF3 was not tested because it does not return a distogram or coordinates for masked residues. Of the complexes for which the sequence-provided prediction was acceptable, 56-65% could be predicted well without use of the peptide sequence (Figure 5c). Examples are shown in Figure 5d to illustrate contact maps and backbone predictions at different RMSD_map_ and scaled peptide RMSD values. The fraction of successes is even greater when shuffling the peptide residues instead of masking them, indicating that amino-acid composition influences the prediction. Performance degrades, however, when using glycines instead of mask tokens. For AF2-Multimer, we did not find a relationship between presence in the training set and the ability of models to make a successful masked peptide prediction (Figure S10a). For Boltz-1 and Chai-1, successful masked predictions had significantly higher numbers of matches in the training set (Figure S10b).

To investigate how the target MSA guides docking for AF2-Multimer, we examined whether sequence conservation explains how models choose binding sites. We annotated the conservation of positions in the target MSA using Jensen-Shannon divergence and observed that binding-site residues tended to be more conserved than non-binding-site residues (Figure S11a). However, predicted binding-site residues and binding-site residues in the native structures had very similar average conservation scores (Figure S11b). Specifically, highly conserved sites were not chosen significantly more frequently than the correct binding site for poor predictions. Figure S11c shows an example where the predicted site is more highly conserved than the native binding site (AF2-Multimer for 3ICI_AC) and a counterexample where the predicted site is much less conserved (Boltz-1 for 6V63_AY) [58, 59, 60, 61]. We also tested whether the models chose binding sites by locating hydrophobic patches on target structures, particularly in cases where the prediction is poor. However, incorrectly predicted binding sites did not have higher average hydrophobicity than native binding sites (Figure S12).

For AF2, the MSA has been hypothesized to aid in an initial global search for the fold, while the structure module further refines the prediction. Consistent with this, in the absence of an MSA, providing a good template rescues performance [62]. To test whether good templates can compensate for the lack of an MSA in protein-peptide docking, we focused on AF2-Multimer and AF3, because Boltz-1 and Chai-1 did not support custom templates at the time of this study. Both AF2-Multimer and AF3 succeed at rigid-body docking when the native structures of the peptide and target are provided with no MSA (see Methods, Figure S13a). For AF3, using native templates modestly outperformed using MSAs without a template (Figure S13a). We found that providing the native template in addition to the target MSA boosted structure prediction accuracy of the target chain and overall docking accuracy (Figure S13b).

To determine whether the target MSA primarily supports target structure prediction or if the evolutionary statistics in the MSA also inform docking, we compared predictions made with the target MSA to predictions where only the target template was provided. AF2-Multimer was sensitive to this difference, performing significantly worse when provided with only the target template, even though the target structure was predicted more accurately (Figure S14a). AF3 performed similarly with both input conditions. We repeated this experiment for the peptide MSA, using the target MSA for the target input. For the subset of complexes where the peptide MSA had at least 50 sequences, we used either the peptide MSA or the native peptide template as the input. AF2-Multimer and AF3 performed slightly better when provided the peptide template (Figure S14b). Most DockQ improvements were small, indicating local structural refinements. These results show that for AF2-Multimer, the evolutionary information available in the target MSA cannot be replaced by structural information, while the evolutionary information in the peptide MSA can be replaced and even improved upon with structural information. To validate this finding further for the target MSA, we repeated the test with masked peptide predictions and, as expected, found that AF2-Multimer performed significantly worse with the target template than with the target MSA, as assessed by RMSD_map_ (Figure S14c). This highlights the importance of the target MSA for AF2 binding site identification.

In most use cases, the bound peptide conformation is not known *a priori*. Thus, we dissected how the peptide MSA is used. We tested whether the peptide MSA contributes to docking performance by removing it for complexes where the peptide MSA (made with default methods) had at least 50 sequences. Interestingly, for most complexes, ablating the peptide MSA had little impact. Overall, peptide MSA ablation slightly reduced model performance, with median DockQ dropping from 0.79-0.85 to 0.69-0.81 across the models (Figure 6a). AF2-Multimer was the most sensitive to this ablation: for complexes for which the original prediction had medium DockQ, 76% maintained medium DockQ after peptide MSA ablation. AF3 was the least sensitive, with 85% of medium predictions maintaining medium DockQ after the ablation. Ablations impacted both the binding site selection and peptide conformation. This can be seen in examples where DockQ dropped from excellent to medium (the peptide conformation was lost) and excellent or medium to poor (the binding site was lost).

**Figure 6.**
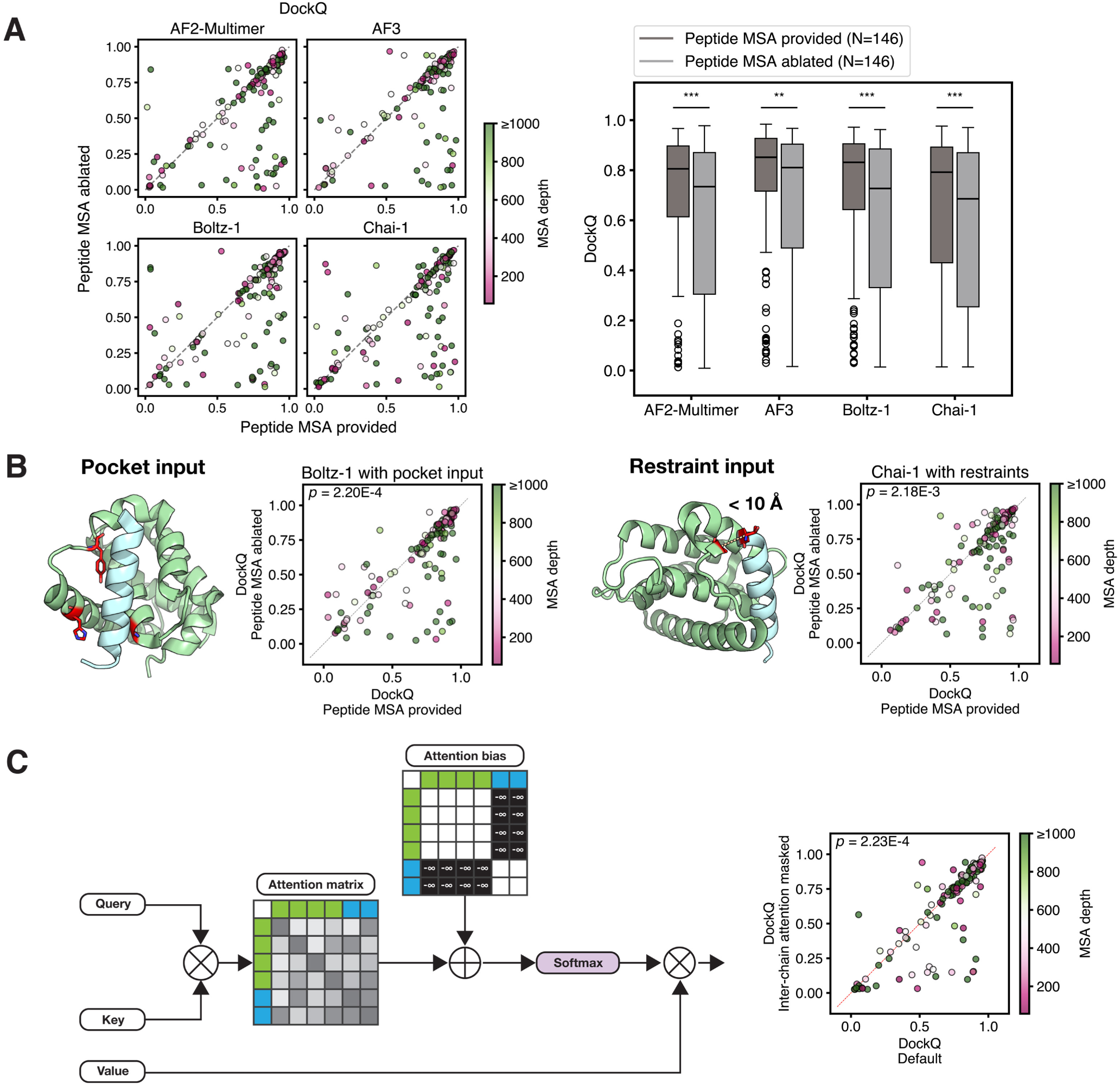
Peptide MSAs boost performance. **(a)** Comparison of DockQ between using the unpaired MSA vs. the peptide-ablated MSA for complexes where the peptide MSA had at least 50 sequences (\**p* < 0.05, \*\**p* < 0.01, \*\*\**p* < 0.001). **(b)** leftmost: Example of Boltz-1 prediction with pocket input, where the three residues in red were designated as pocket residues; middle left: Comparison of DockQ scores using the unpaired MSA vs. peptide-ablated MSA when Boltz-1 pocket input was provided; middle right: Example of Chai-1 prediction with restraint input, where the interaction between the two residues was provided as a 10 Å distance restraint; right: Comparison of DockQ scores using the unpaired MSA vs. peptide-ablated MSA when Chai-1 restraint input was provided. **(c)** left: Procedure for masking interchain attention in the Evoformer module of AF2-Multimer; right: DockQ for predictions made with inter-chain attention unmasked (horizontal axis) vs. inter-chain attention masked (vertical axis). All *p*-values are from a Wilcoxon signed-rank test comparing the two distributions.

To assess the extent to which the peptide MSAs inform the peptide binding pose, beyond binding site prediction, we made predictions where the binding site was provided to the model. We focused on Boltz-1 and Chai-1 for this experiment because these models provide the option to specify binding-site residues or distance restraints. For Boltz-1, three binding-site residues from the target were provided to the model as part of the “pocket” (Figure 6b, left). For Chai-1, we included a single distance restraint between a binding-site residue and a peptide residue (*≤* 10 Å) (Figure 6b, right). Providing constraints improved model accuracy (Figure S15). But when constraints were provided, ablating the peptide MSA (for peptide MSAs with at least 50 sequences) still resulted in a decrease in model accuracy, demonstrating that the peptide MSA contributes to peptide pose prediction (Figure 6b).

To understand how AF2-Multimer processes unpaired protein and peptide MSA information together in the Evoformer module, we masked row- and column-wise self-attention between rows and columns corresponding to different chains in the MSA (Figure 6c, left). In this masked attention regime, the model may leverage intra-chain evolutionary information and develop an understanding of each chain individually, but cannot process the two MSAs together in the Evoformer module. We saw significantly lower performance with this masking protocol on some of the complexes where the peptide MSA had at least 50 sequences (Figure 6c, right). Although coevolutionary statistics cannot be directly extracted, as is possible for a paired MSA, the Evoformer is nonetheless able to learn from inter-chain attention on an unpaired MSA.

### Explaining successful AF3 predictions of novel complexes

All models perform very well on complexes with homologs in the training data, suggesting that successful prediction may be achieved using a kind of homology modeling. Nevertheless, some test cases that lacked similarity to training-set complexes were still accurately predicted. We manually investigated the six novel complexes that AF3 predicted with DockQ > 0.49: 8BRH_AB, 8S6O_DH, 8FG6_BA, 8HDJ_BA, 8TEE_AC, and 9G6Z_AB [43, 63, 64, 65, 66, 67, 68, 69, 70, 71, 72]. For 8BRH_AB, AF3 was successful when the target MSA was provided but unsuccessful when only the target template was provided (Figure S16a). We conclude that there is some essential evolutionary information present in the target MSA and note that the native binding site is more highly conserved than the rest of the target, which could facilitate binding site identification. In contrast, for 8S6O_DH, AF3 was able to dock the peptide when provided the target template and no MSA (Figure S16b, upper left). Since the native binding site is a hydrophobic cleft and the peptide has two aromatic residues, we hypothesized that AF3 chooses a pose that buries hydrophobic residues (Figure S16b, upper right). Supporting this, when Phe and Tyr in the peptide DE**F**I**Y**P were mutated to alanine, AF3 did not recover the native peptide backbone conformation but still chose the native binding site (Figure S16b, lower left). When all residues in the target binding site were mutated to alanine, AF3 no longer predicted that site (Figure S16b, lower right). For the remaining four examples, we attribute the success of AF3 to the identification of common secondary structure packing motifs across the interface. For example, for 8FG6_BA and 8HDJ_BA, AF3 consistently captured the *β*-sheet formed by the peptide and the correct binding site (Figure S16c, first and second row). For 8TEE_AC, the native structure also features a *β*-sheet interaction. When the 12 interfacial peptide residues were mutated to alanines, which have high helical propensity, AF3 predicted a helical conformation for the peptide and repositioned it to pack against an *α*-helix in the target (Figure S16c, third row). Here, the predicted secondary structure of the peptide influenced the binding mode. Finally, with 9G6Z_AB, AF3 predicted the peptide to adopt an *α*-helix at the native binding site, which consists of several small helices (Figure S16c, last row).

## Discussion

In this study, we evaluated the performance of AF2-Multimer, AF3, Boltz-1, and Chai-1 on the task of protein-peptide complex structure prediction. Most predictions identified the correct binding site, with DockQ > 0.49, and many were highly accurate. AF3 slightly outperformed Boltz-1 and Chai-1, potentially due to a larger distilled training data set or increased ability to memorize information in the training set, as indicated by the sensitivity of the performance of AF3 to training set similarity [19, 20]. Data leakage into test sets constructed based on a training-date cutoff is likely for the PDB, where many proteins and protein complexes are represented with hundreds of related, although non-identical, structures. To address this, we added another test for training-test set overlap based on complex structural similarity. For all models, binding sites that were more represented in the training set were predicted more accurately. In some cases, the training data may have led the model to use an incorrect but commonly seen binding site. Of incorrect predictions with DockQ < 0.23, 79-83% involved complexes with one or more characteristics correlated with poorer performance, including irregular secondary structure or binding-site matches to an incorrect site in the PDB (Figure S17).

Our findings align with a previous benchmark of AF2-Multimer reporting that the model struggles with predicting PROTAC-mediated protein-protein interfaces [73], which tend to involve proteins that do not interact in natural systems. Additionally, our observations align with benchmarking of AF3, Boltz-1, and Chai-1 on protein-ligand binding, where ligands with abundant data in the PDB show better prediction performance [74]. It is still common practice to use homology splits based on single chains and/or temporal splits for training and evaluation. The large differences in prediction quality for pre- vs. post-training examples and examples with vs. without homologous matches in the training data highlight the need for non-redundant evaluation sets to estimate performance on novel tasks. Bias for previously observed complexes raises a concern for generalization to novel interactions and targeting new binding sites when designing peptide binders. Prediction tasks of increasing complexity, e.g., for protein-protein interactions and biomolecular assemblies that are not well-represented in the limited PDB, also require this type of extrapolation.

Although previously seen structures are predicted better, we found cases where novel interactions were predicted correctly. Additionally, there were cases of PDB structures present in the training set that the models failed to predict. These surprising examples led us to investigate the role of the MSA input. First, we found that models do not rely on inter-chain coevolutionary signal. This is in contrast with the predominant theory that AF2-Multimer folds protein-protein complexes based on coevolutionary information between chains, which led the developers of AF2 and other models to pair sequences by species in the MSA construction process. Further, for protein-protein interactions, Bryant et al. found that a paired + unpaired MSA resulted in the best predictions for AF2 with monomer-trained weights and that the number of sequences in the paired MSA has a strong influence on predictions [13]. These observations from the literature do not clarify whether the pairing in the MSA or the increased depth of the MSAs is responsible for the boost in prediction quality. In our experiments, we found weak evidence that models have learned to incorporate coevolution (as defined by patterns detectable after sequence pairing) into predictions of protein-protein or protein-peptide complexes.

In the absence of a paired MSA, we found that the target MSA contains sufficient information for the prediction of binding sites in many cases and that this information is not just used to inform the structure of the target. Conserved positions on the target do not, alone, explain the choice of binding site, though previous work in tuning the AF2 MSA profile for improving predictions suggests that statistics of residue frequencies provided to the model play an important role [75, 76]. Across the models, we found that the peptide MSA improves 15-24% of complex predictions, influencing both binding-site and binding-pose prediction. With AF2-Multimer, we found evidence indicating that inter-chain interactions are implicitly extracted from an unpaired MSA through the Evoformer mechanism of interleaving row- and column-wise attention. This is a previously undescribed mechanism for how AF2-Multimer treats unpaired MSAs. Because the MSA modules of AF3 and related models do not use column-wise attention, we leave the interrogation of how inter-chain statistics are processed in these models to future work.

Close inspection of the few highly successful novel predictions by AF3 showed that most recognize common secondary structure element interactions at the interface. We found one example where the evolutionary information in the target MSA was critical for docking success. While binding site conservation and hydrophobicity are possible explanations for the accurate modeling of 8BRH_AB and 8S6O_DH, respectively (Figure S16a), these were not strongly predictive features in our benchmark overall. Our results demonstrate that peptide docking success cannot be explained using simple MSA-based statistics alone, since models have learned to rely on sources of information beyond covariation. Studies aimed at explaining the behavior of protein language models, such as ESM, have been fruitful [77, 78, 79, 80]. Similar efforts for MSA-based sequence-to-structure models will continue to advance our understanding of how these widely used methods make predictions, particularly as MSA subsampling and tuning have brought new capabilities to structure prediction and design methods [81, 75, 76, 82].

---

**Figure S1.**
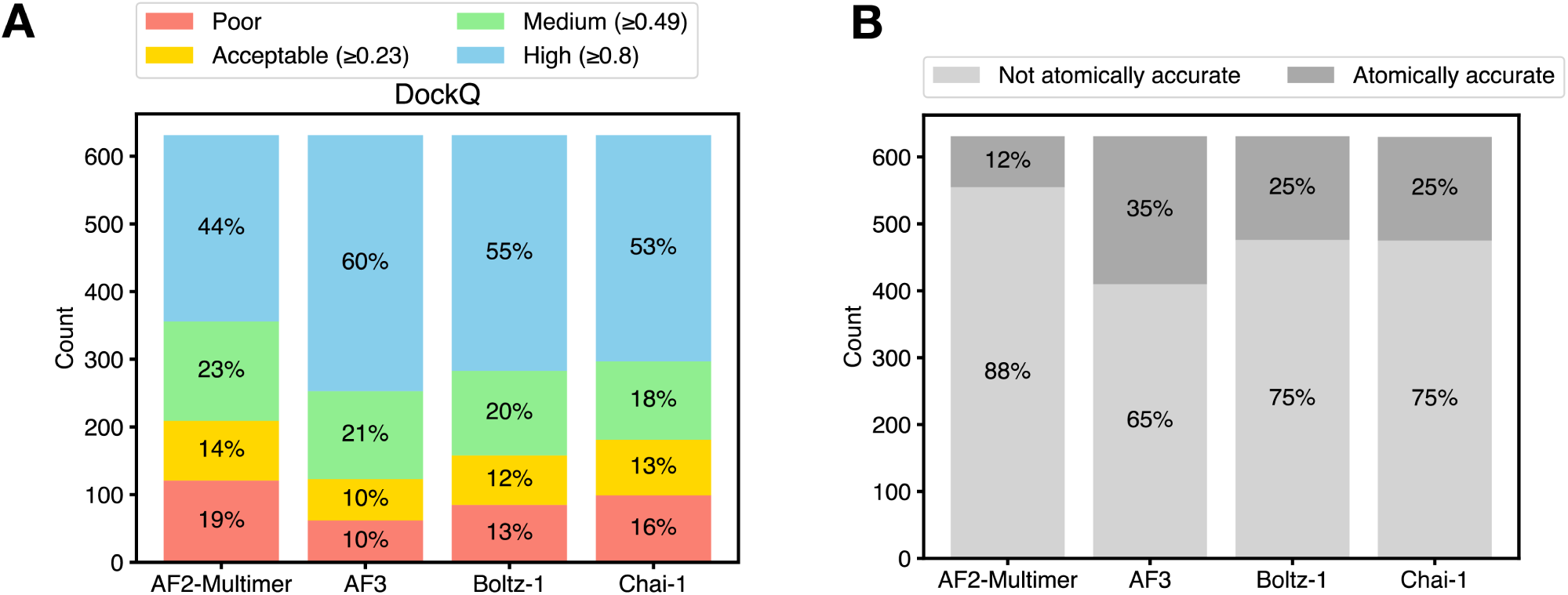
Summary of docking accuracy. **(a)** Fraction of predictions in each DockQ bin, using categories from the CAPRI assessment [1, 2, 3]. **(b)** Fraction of predictions that are atomically accurate, as defined by the following criteria: >90% of native contacts recovered (contacts within 4 Å), no clashes, peptide all-atom RMSD < 2 Å, interface all-atom RMSD < 3 Å (see Methods).

**Figure S2.**
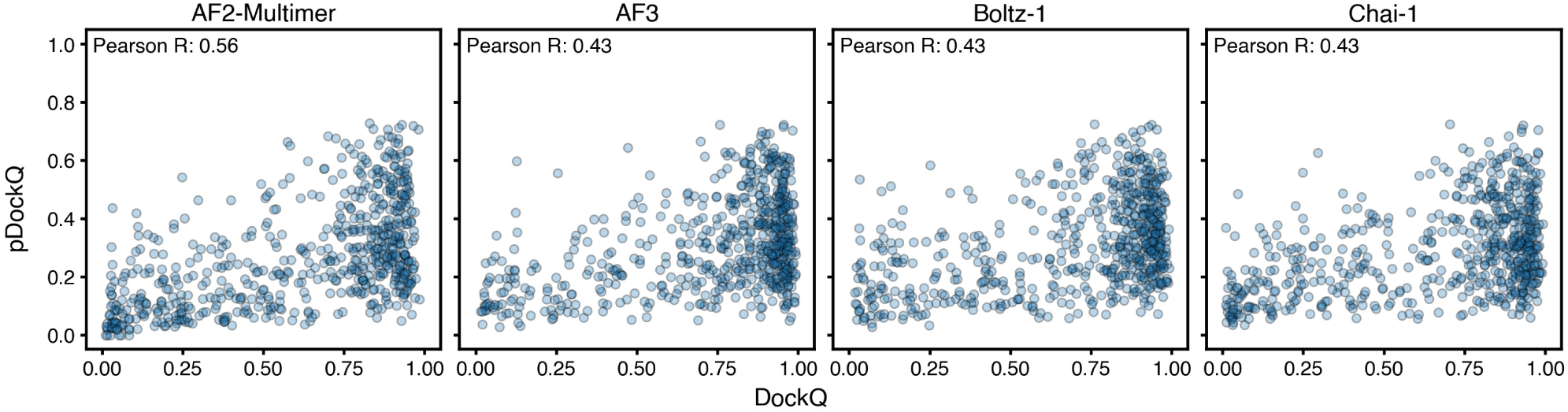
Model confidence and accuracy as measured by pDockQ and DockQ.

**Figure S3.**
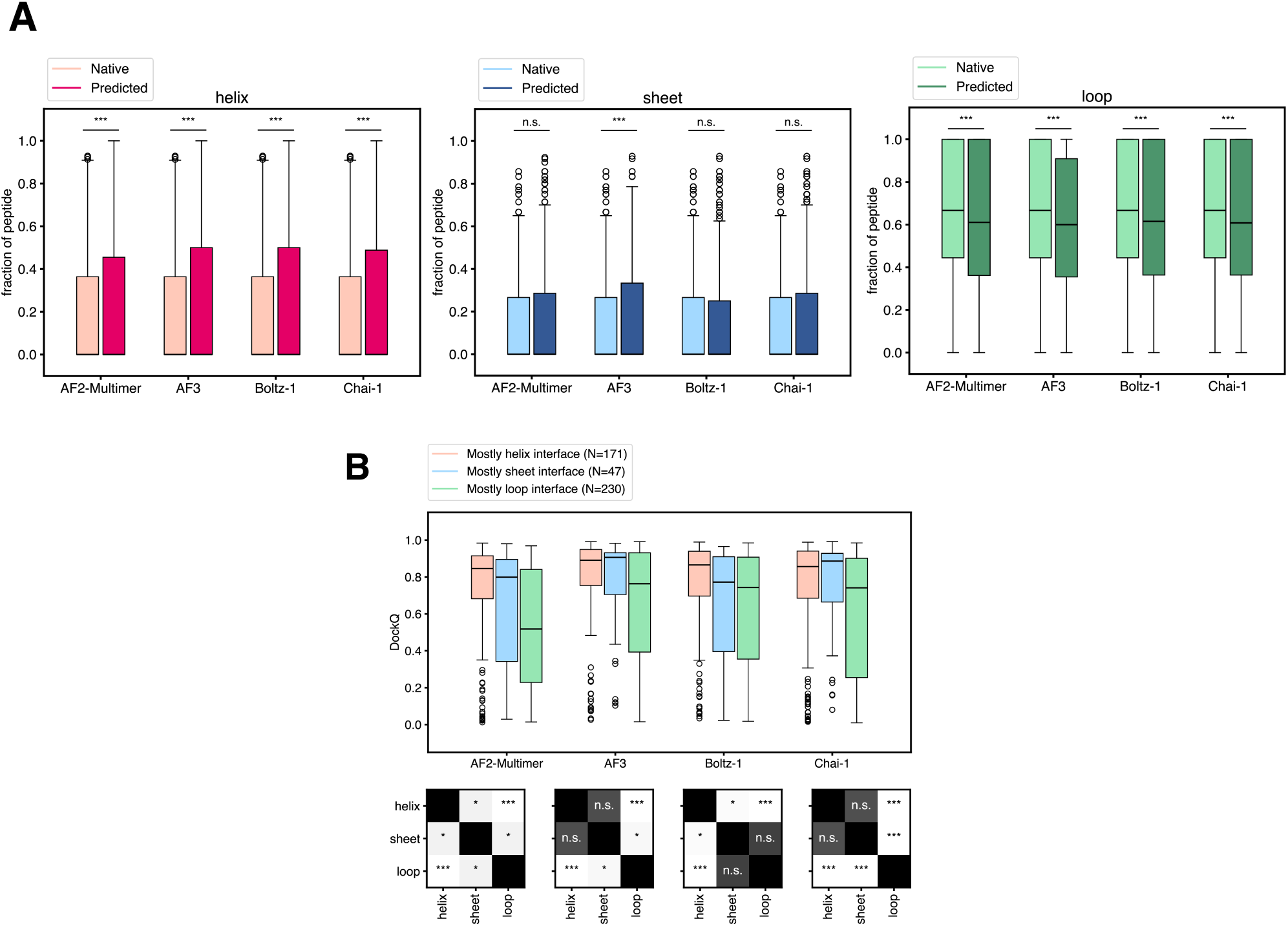
Secondary structure biases in predicted complex structures. **(a)** Secondary structure composition of native peptides and peptide structures predicted by the different models. The fraction of peptide residues adopting a certain secondary structure is shown. (Wilcoxon signed-rank test, \**p* < 0.05, \*\**p* < 0.01, \*\*\**p* < 0.001). **(b)** Distribution of DockQ for complexes where the native interface had >50% residues with helical, sheet, or loop-like secondary structure. *p*-values shown in table: Mann-Whitney U-test, \**p* < 0.05, \*\**p* < 0.01, \*\*\**p* < 0.001.

**Figure S4.**
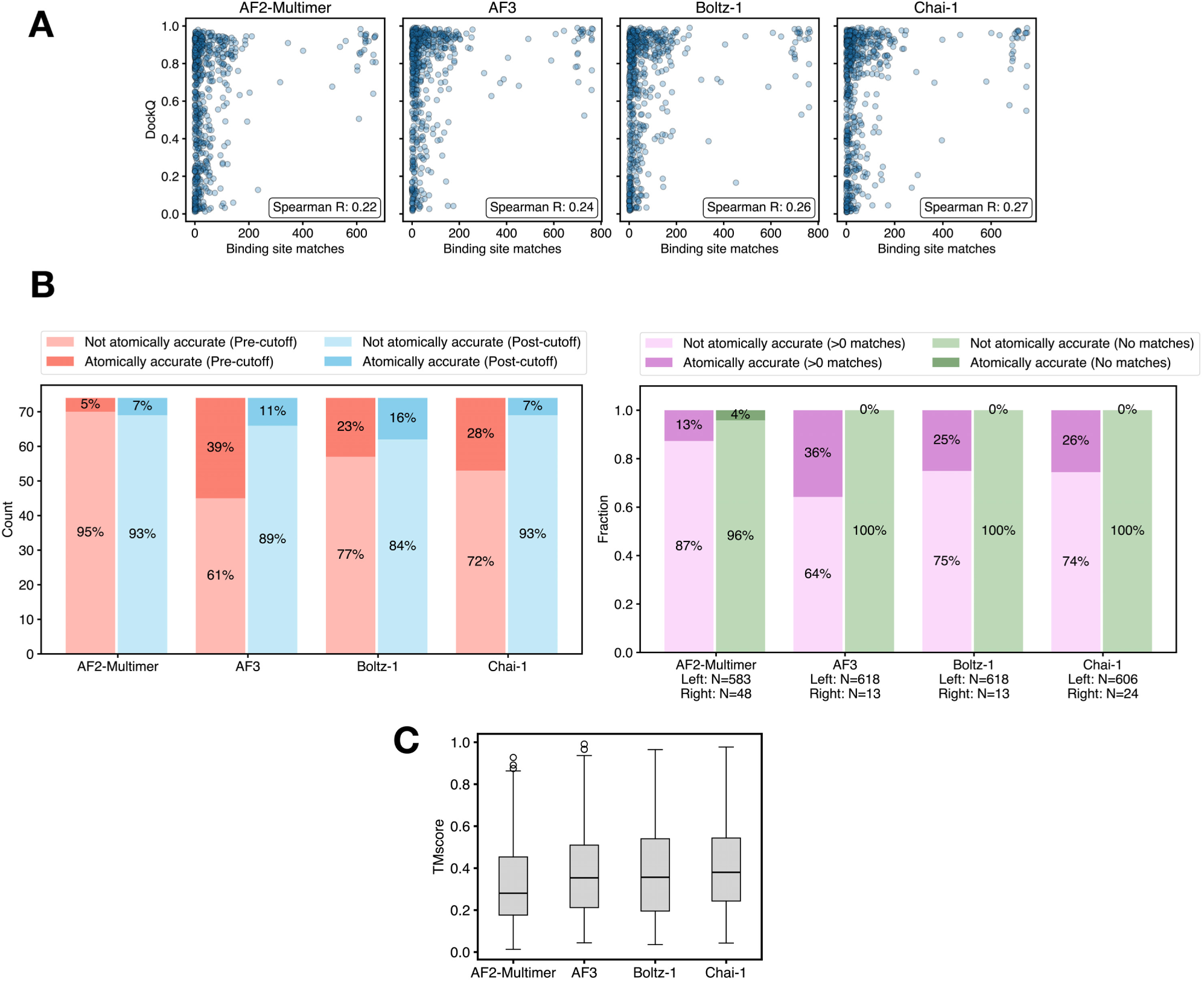
Prediction accuracy is higher for structures with similarity to the training set. **(a)** DockQ and number of binding site matches. **(b)** Fraction of atomically accurate predictions for the (left) pre-cutoff and post-cutoff sets, N=74 for each and (right) the > 0 training set matches and 0 training set matches sets. Atomic accuracy is defined by the following criteria: >90% of native contacts recovered (contacts within 4 Å), no clashes, peptide all-atom RMSD < 2 Å, interface all-atom RMSD < 3 Å (see Methods). **(c)** Distribution of TMscores between the peptide predicted alone and the native bound peptide structure.

**Figure S5.**
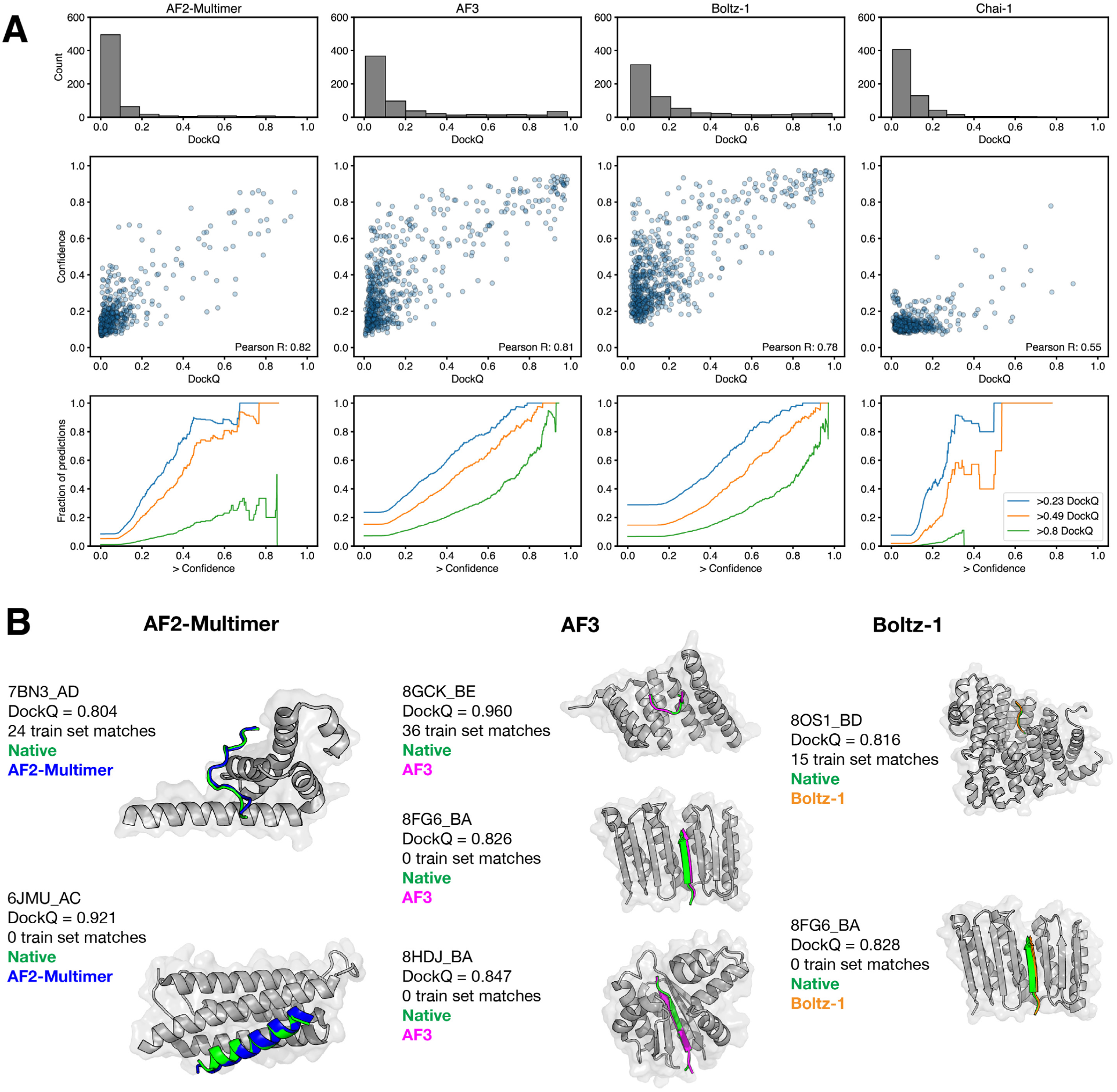
Predictions are rarely accurate without an MSA. **(a)** Model confidence and accuracy when no MSA is provided for either chain. **(b)** Complexes where MSA-ablated predictions have DockQ > 0.8 and the structures were released after each model’s respective training set date cutoff. Training set matches were determined through the same procedure used in Figure 2c-d (see Methods).

**Figure S6.**
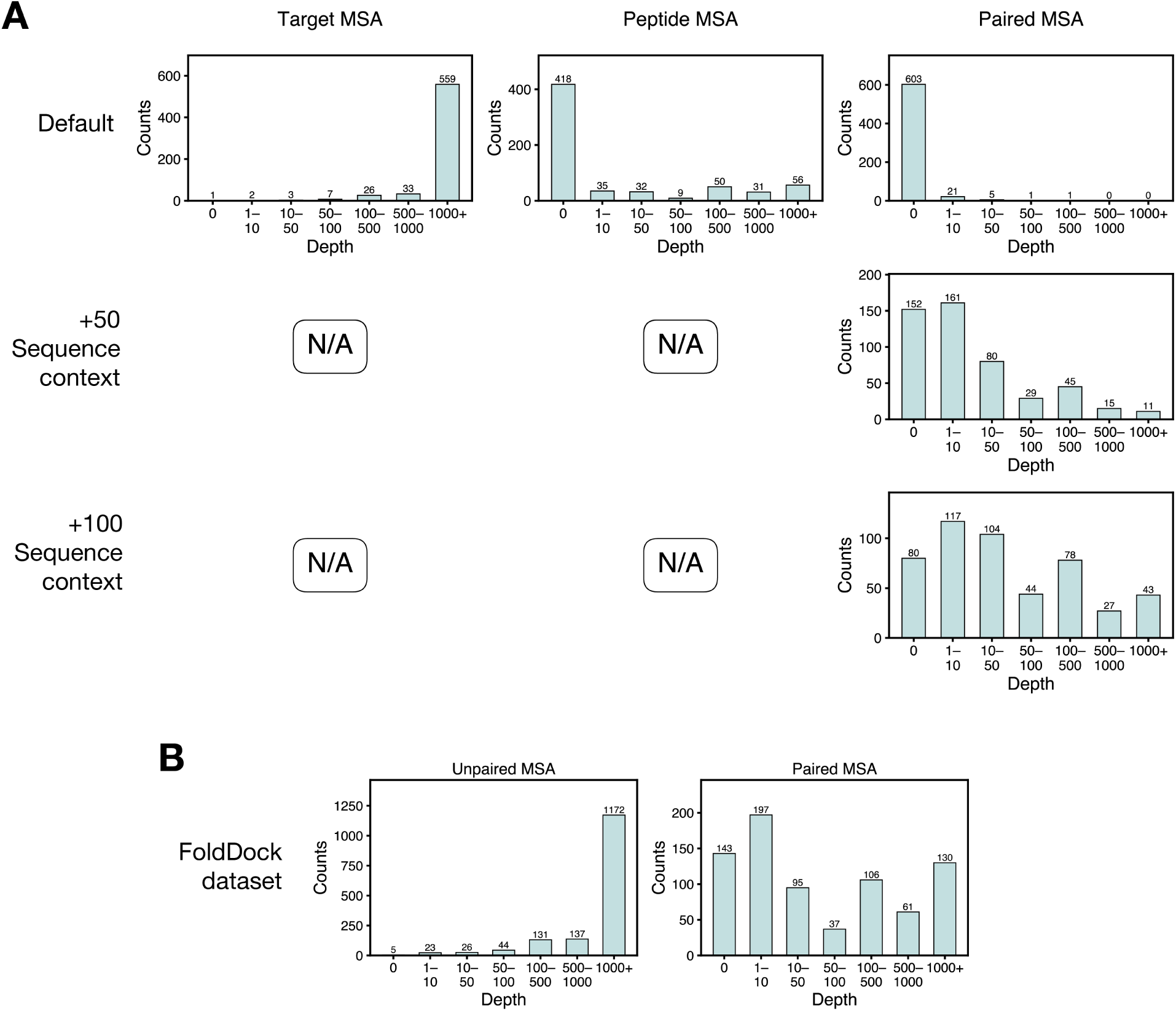
MSA depths for various inputs. **(a)** For Default predictions, target MSA depths and peptide MSA depths correspond to MSAs generated using the mmseqs server where only an unpaired MSA was requested. The paired MSA depths correspond to MSAs generated where only a paired MSA was requested. For +50 and +100 sequence context, depths are shown only for MSAs where the peptide could be mapped to a UniProt protein, though predictions were made on only those containing at least 50 sequences and passing a quality filter. **(b)** MSA depths for the FoldDock PPI dataset. The unpaired MSA depth is reported per-chain, while each paired MSA depth corresponds to the paired MSA of a complex. All MSA depths are shown, though predictions were made on only those containing at least 50 sequences.

**Figure S7.**
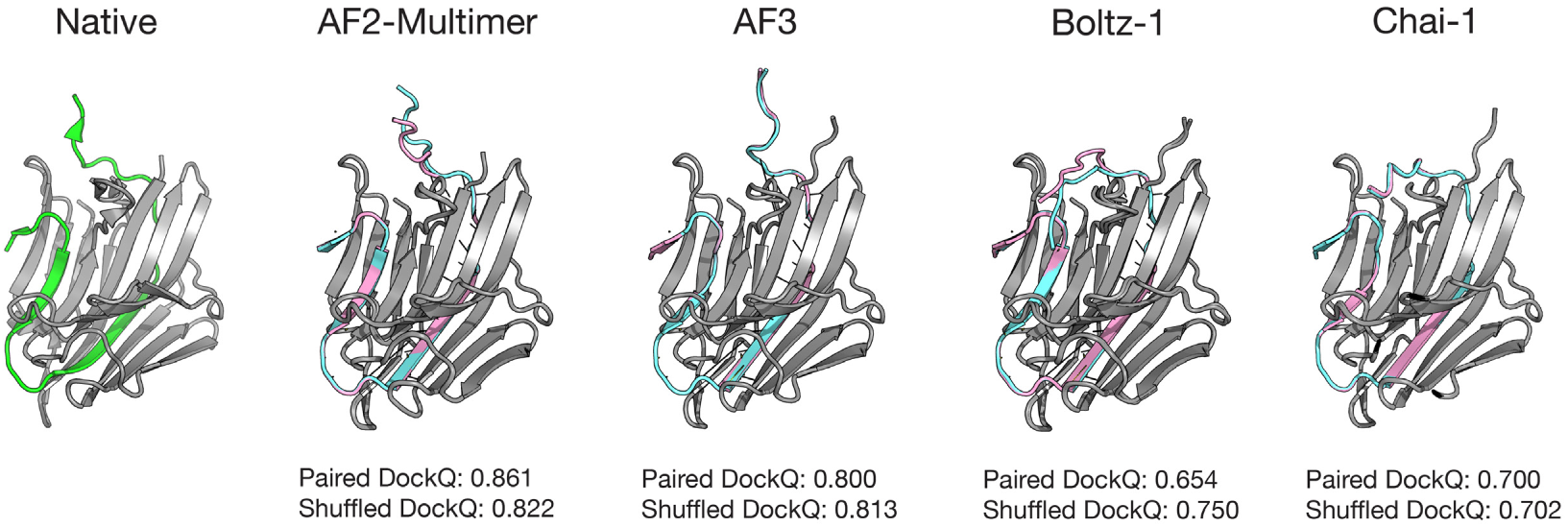
Predictions for 3TRS_BA with +100 context paired MSA (cyan) and +100 context shuffled MSA (pink)

**Figure S8.**
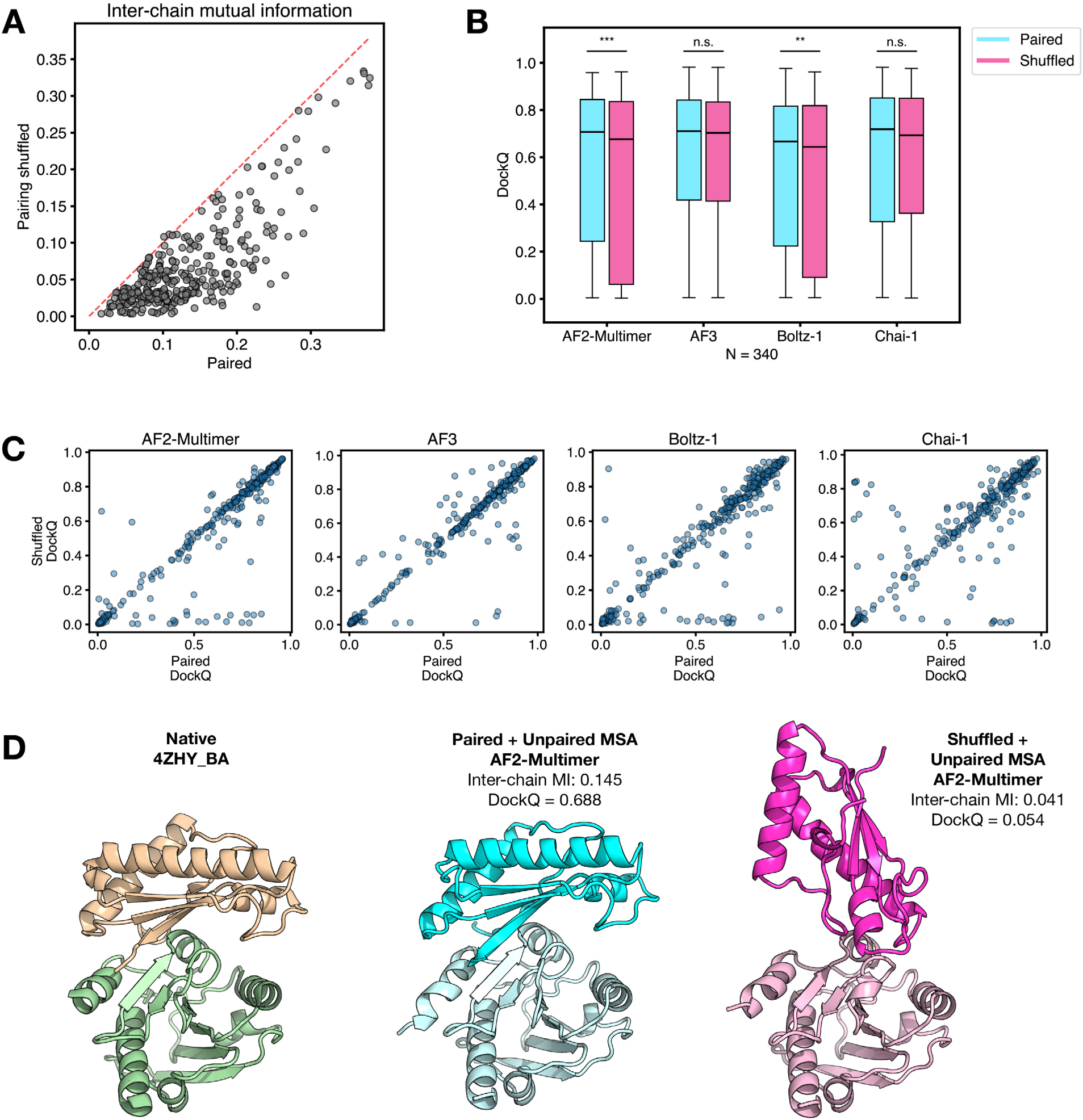
MSA pairing does not improve prediction performance for protein-protein interactions. **(a)** Distribution of MI for predictions made with an unpaired + paired MSA and unpaired + shuffled MSA for protein-protein complexes reported in Bryant et al [4]. Complexes where the paired MSA had depth at least 50 sequences are included. (b) Distribution of DockQ for predictions made with unpaired + paired vs. unpaired + shuffled MSAs (Wilcoxon signed-rank test, \**p* < 0.05, \*\**p* < 0.01, \*\*\**p* < 0.001). (c) Scatter plot of DockQ for predictions made with unpaired + paired vs. unpaired + shuffled MSAs. (d) The native structure and AF2-Multimer predictions for 4ZHY_BA with paired + unpaired MSA and shuffled + unpaired MSA [5, 6].

**Figure S9.**
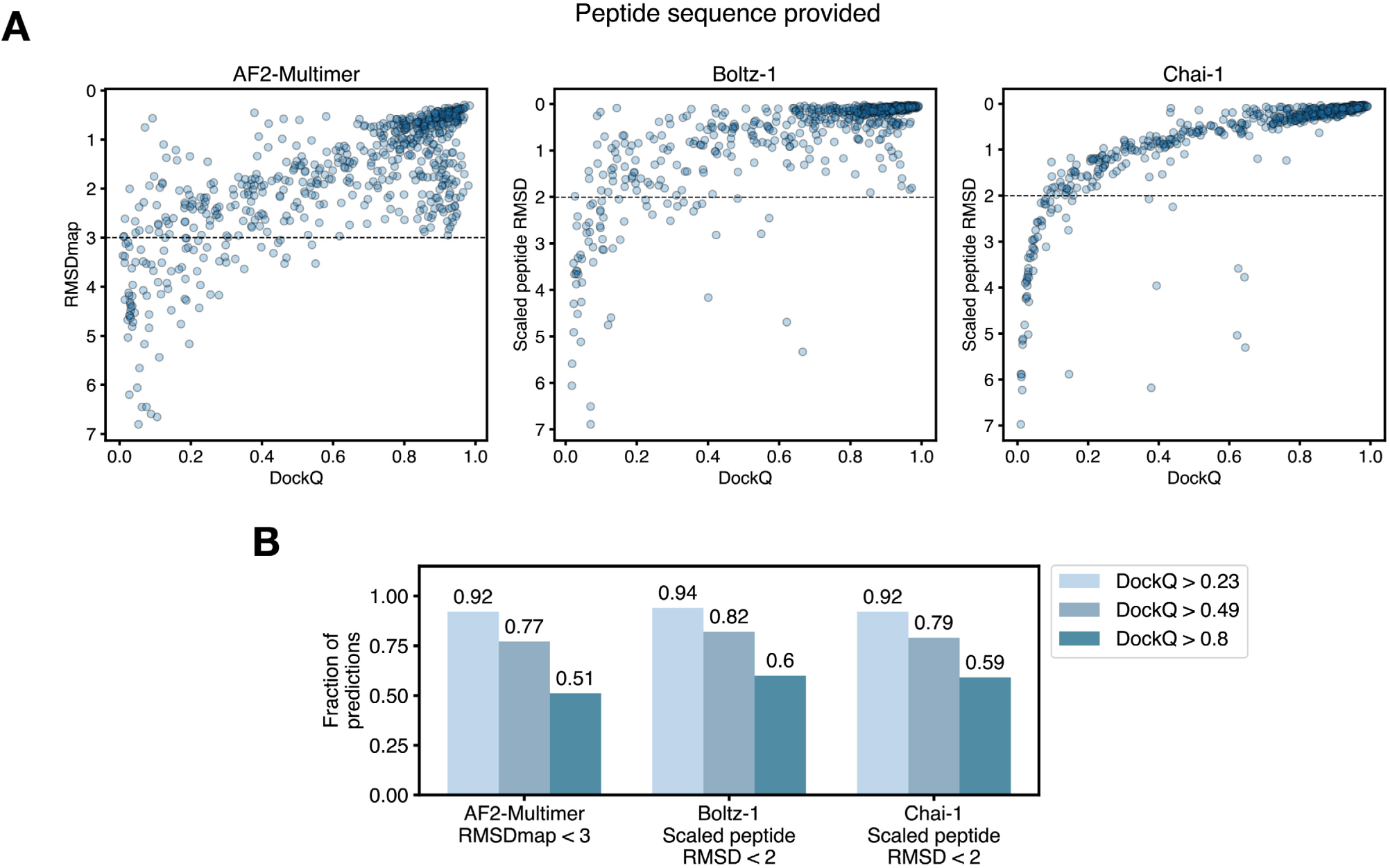
RMSDmap and scaled peptide RMSD correlate with DockQ. **(a)** PPerformance metric vs. DockQ for predictions where the peptide sequence is provided. For AF2-Multimer, the performance metric is contact map RMSD (RMSDmap). For Boltz-1 and Chai-1, the performance metric is scaled peptide RMSD. **(b)** Fraction of predictions with DockQ > *z*, for different *z*, using an RMSDmap cutoff of 3 and with a scaled peptide RMSD cutoff of 2 Å.

**Figure S10.**
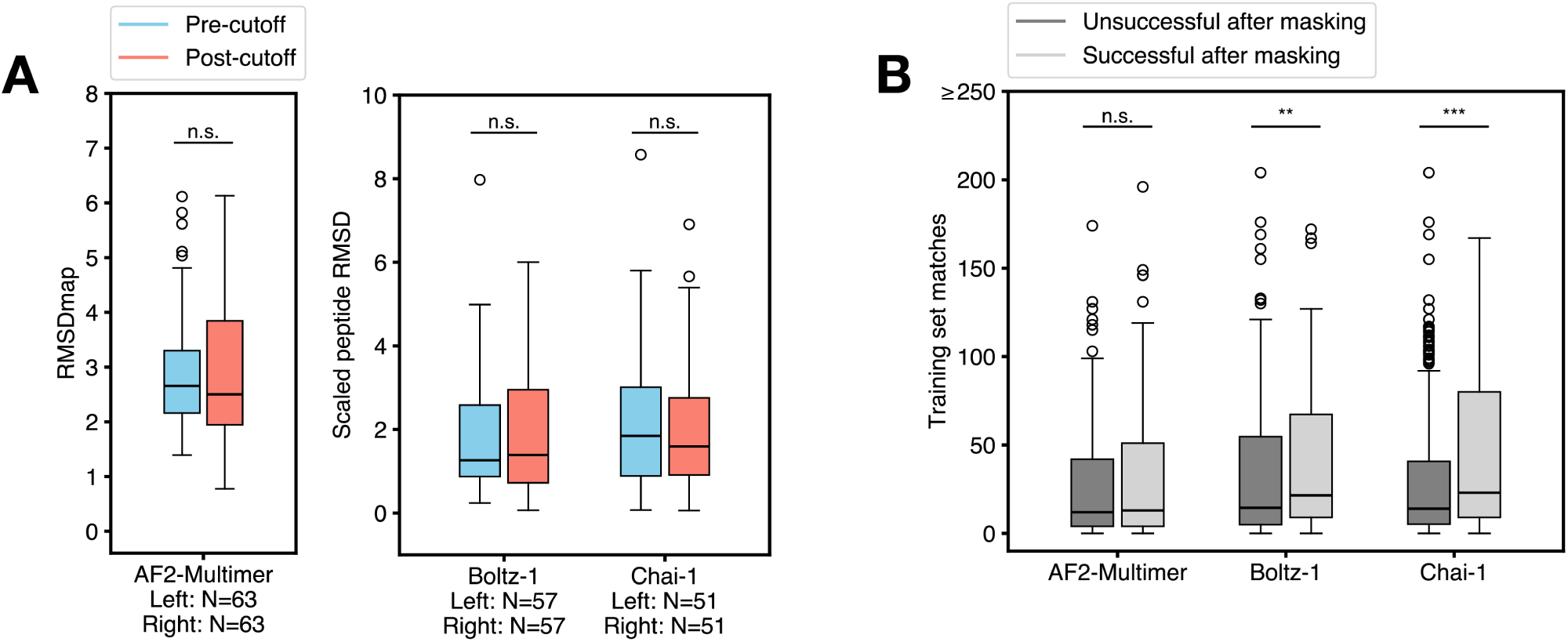
**(a)** RMSDmap and scaled peptide RMSD values for pre-cutoff and post-cutoff masked predictions. Only complexes that were successful unmasked predictions are shown. **(b)** Number of training set matches for complexes that had an unsuccessful vs. successful masked prediction. Only complexes that were successful unmasked predictions are shown.

**Figure S11.**
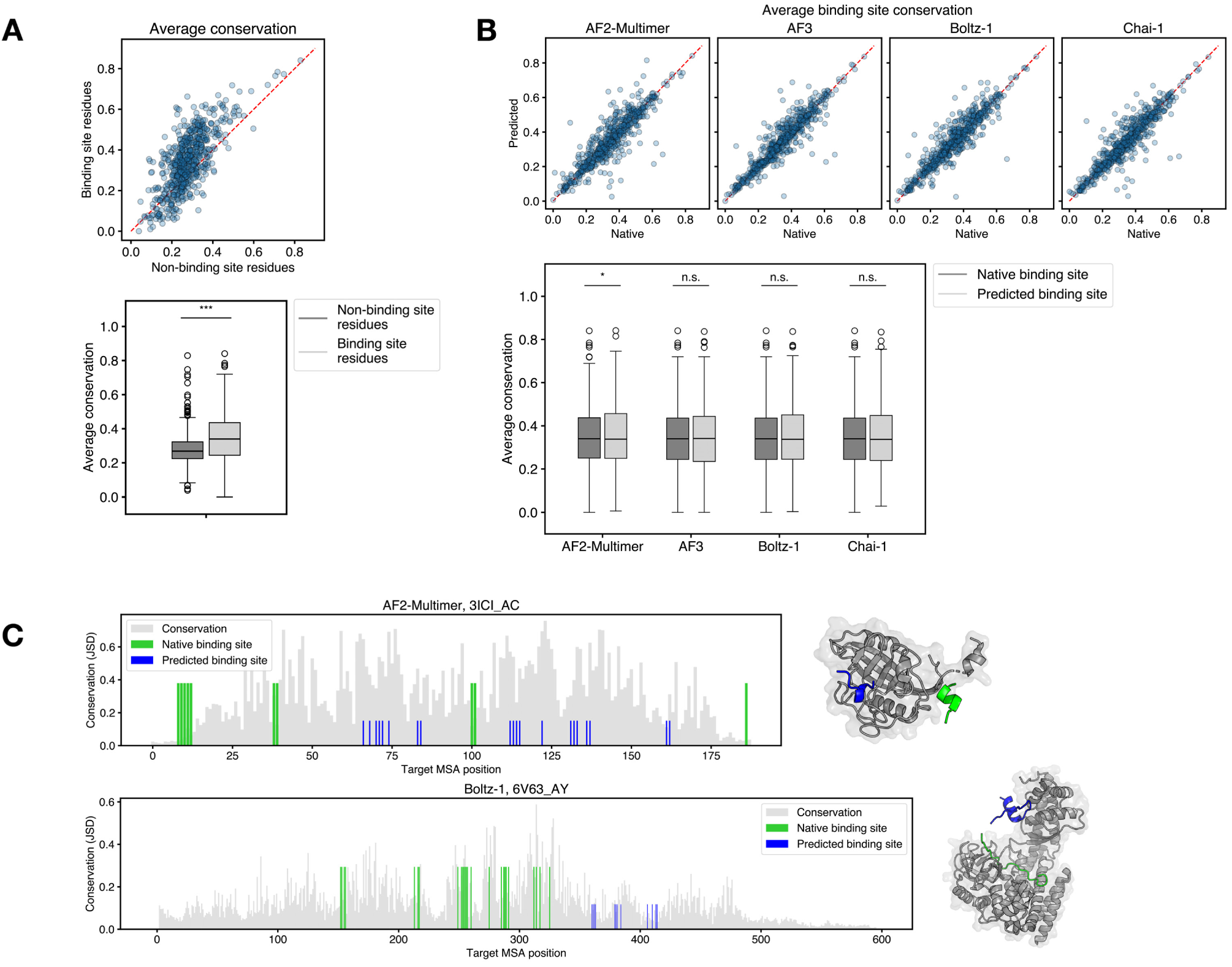
Predicted binding sites are not more conserved than native binding sites. **(a)** Average binding site conservation of native binding site residues vs. native non-binding residues. (Mann-Whitney U-test, \**p* < 0.05, \*\**p* < 0.01, \*\*\**p* < 0.001). **(b)** Binding site conservation of predicted and native structures. All p-values are from a Wilcoxon signed-rank test comparing the two distributions. **(c)** Examples of conservation of binding-site residues on the target for native and predicted structures.

**Figure S12.**
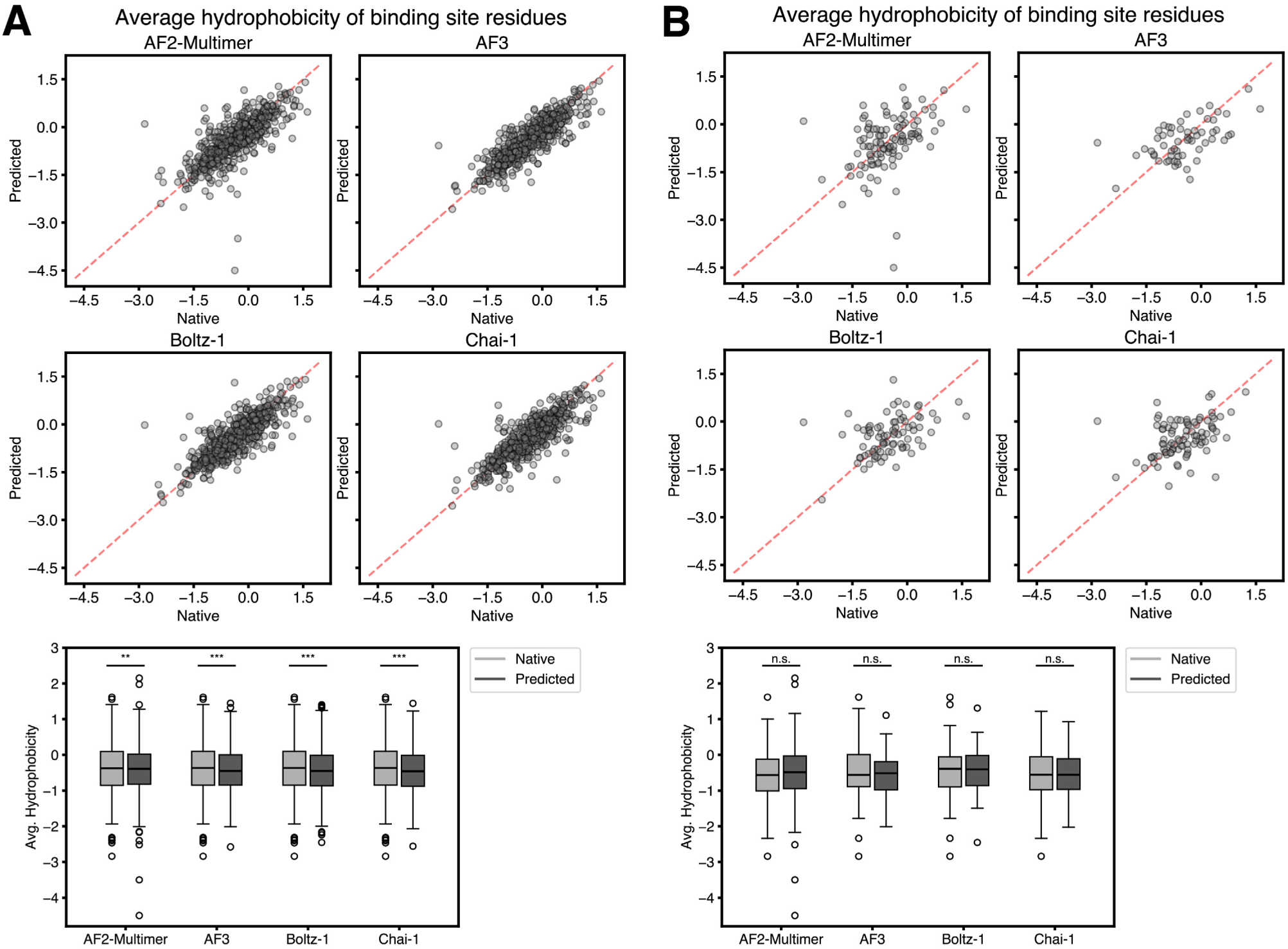
The predicted binding site is not determined by hydrophobicity. **(a)** Average hydrophobicity of binding-site residues in native and predicted structures. **(b)** Average hydrophobicity of binding-site residues in native and predicted structures for predictions of DockQ < 0.23. (Wilcoxon signed-rank test, \**p* < 0.05, \*\**p* < 0.01, \*\*\**p* < 0.001).

**Figure S13.**
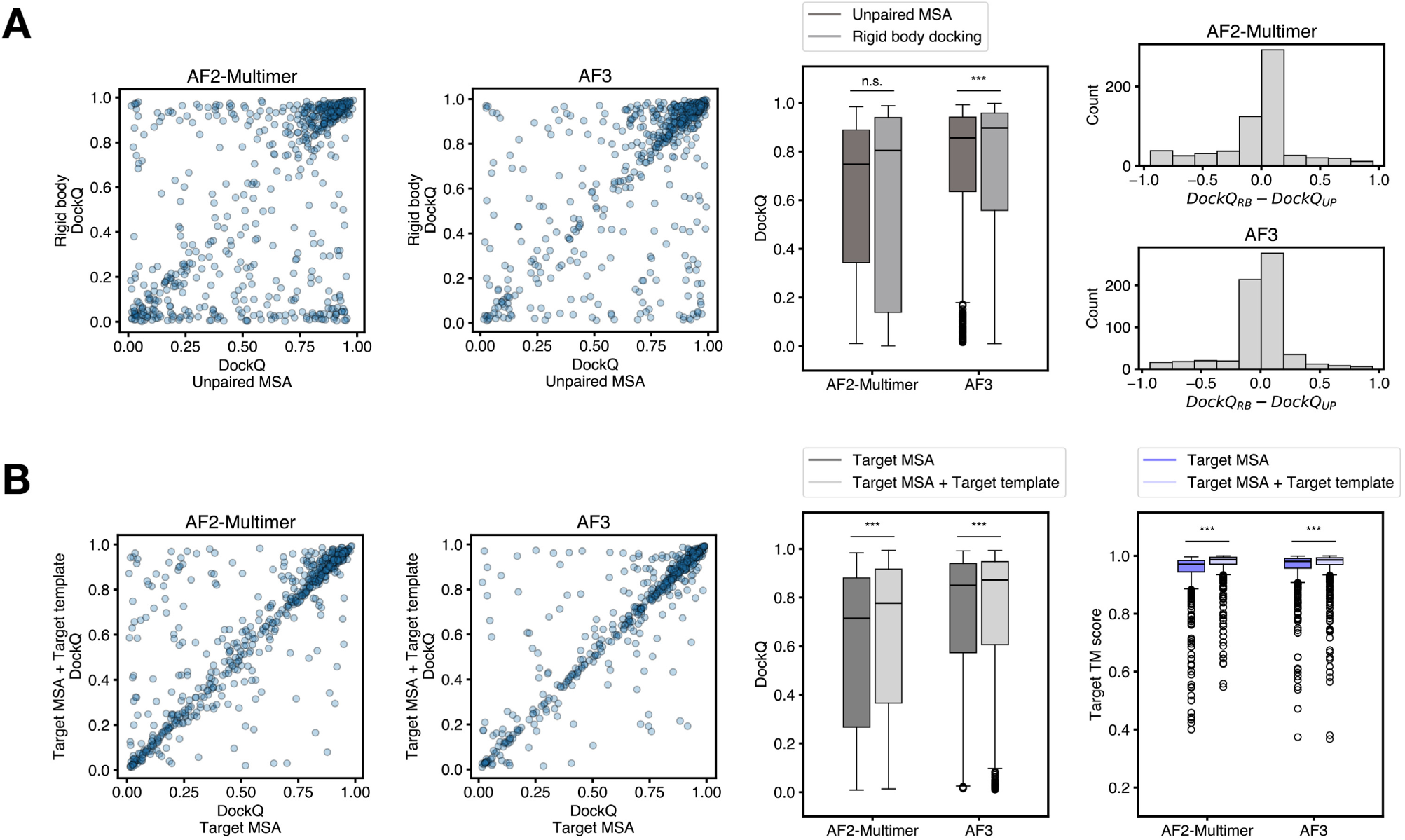
**(a)** (left and middle) DockQ for predictions made with an unpaired MSA or with templates provided for each chain and no MSA (“rigid-body docking”). (right) Difference in DockQ between rigid-body predictions and unpaired MSA predictions for the two models. **(b)** (left and middle) DockQ for predictions made with just the target MSA or with the target MSA + target template. (right) Distributions of Target TM score for the two conditions. (Wilcoxon signed-rank test, \**p* < 0.05, \*\**p* < 0.01, \*\*\**p* < 0.001).

**Figure S14.**
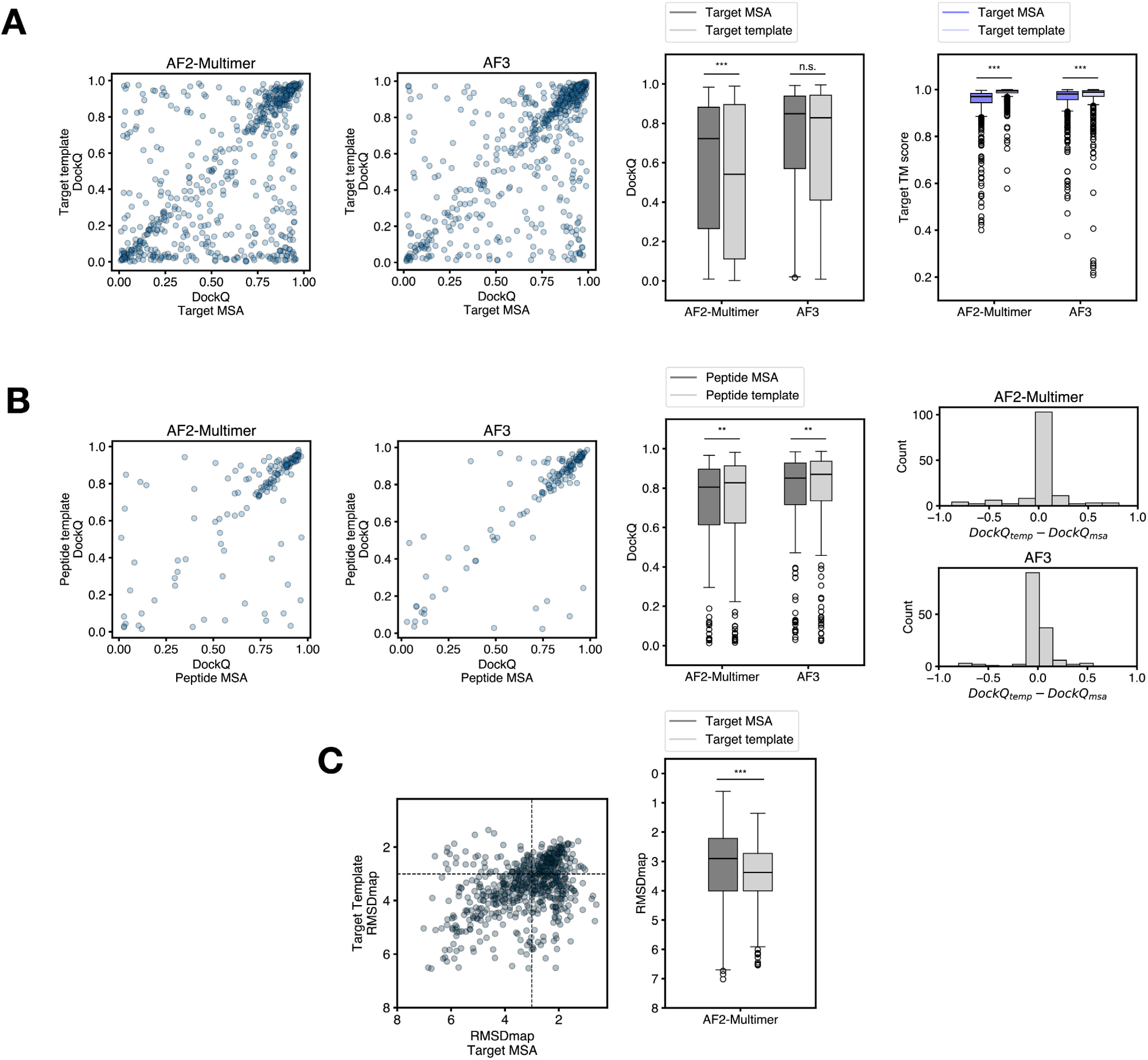
**(a)** left and middle: DockQ for predictions made with just the target MSA or with just the target template as input; right: Distributions of target TM score for the two conditions. **(b)** left and middle: DockQ for predictions made the target MSA + peptide MSA or with the target MSA + peptide template; right: Difference in DockQ between target MSA + peptide template and target MSA + peptide MSA for the two models (Wilcoxon signed-rank test, \**p* < 0.05, \*\**p* < 0.01, \*\*\**p* < 0.001). **(c)** RMSDmap values for masked predictions made with just the target MSA or with just the target template as input.

**Figure S15.**
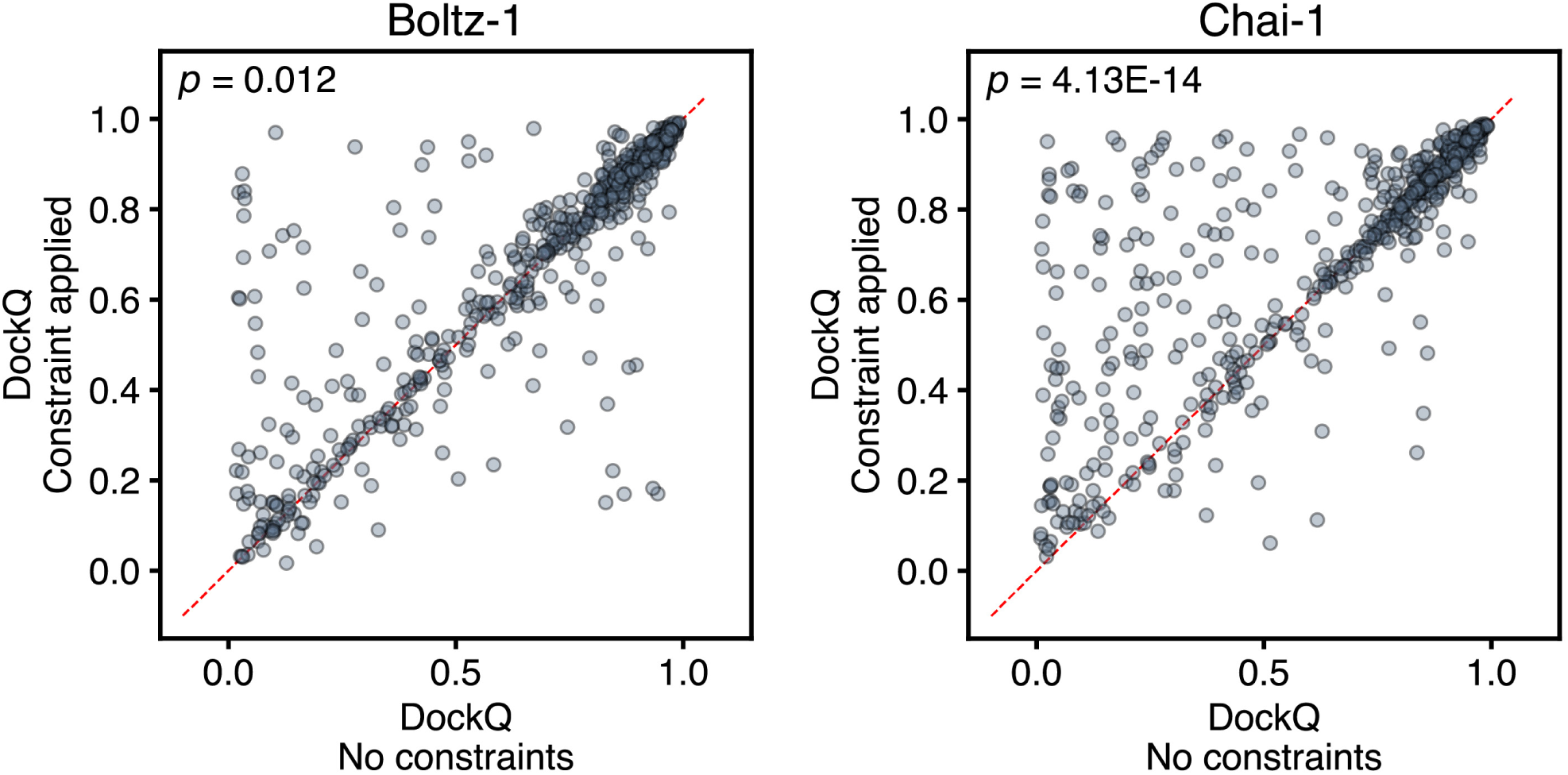
Model accuracy when pocket input for Boltz-1 and restraint input for Chai-1 are provided. *p*-values reflect the difference in DockQ distributions between constrained prediction and unconstrained prediction (Wilcoxon signed-rank test).

**Figure S16.**
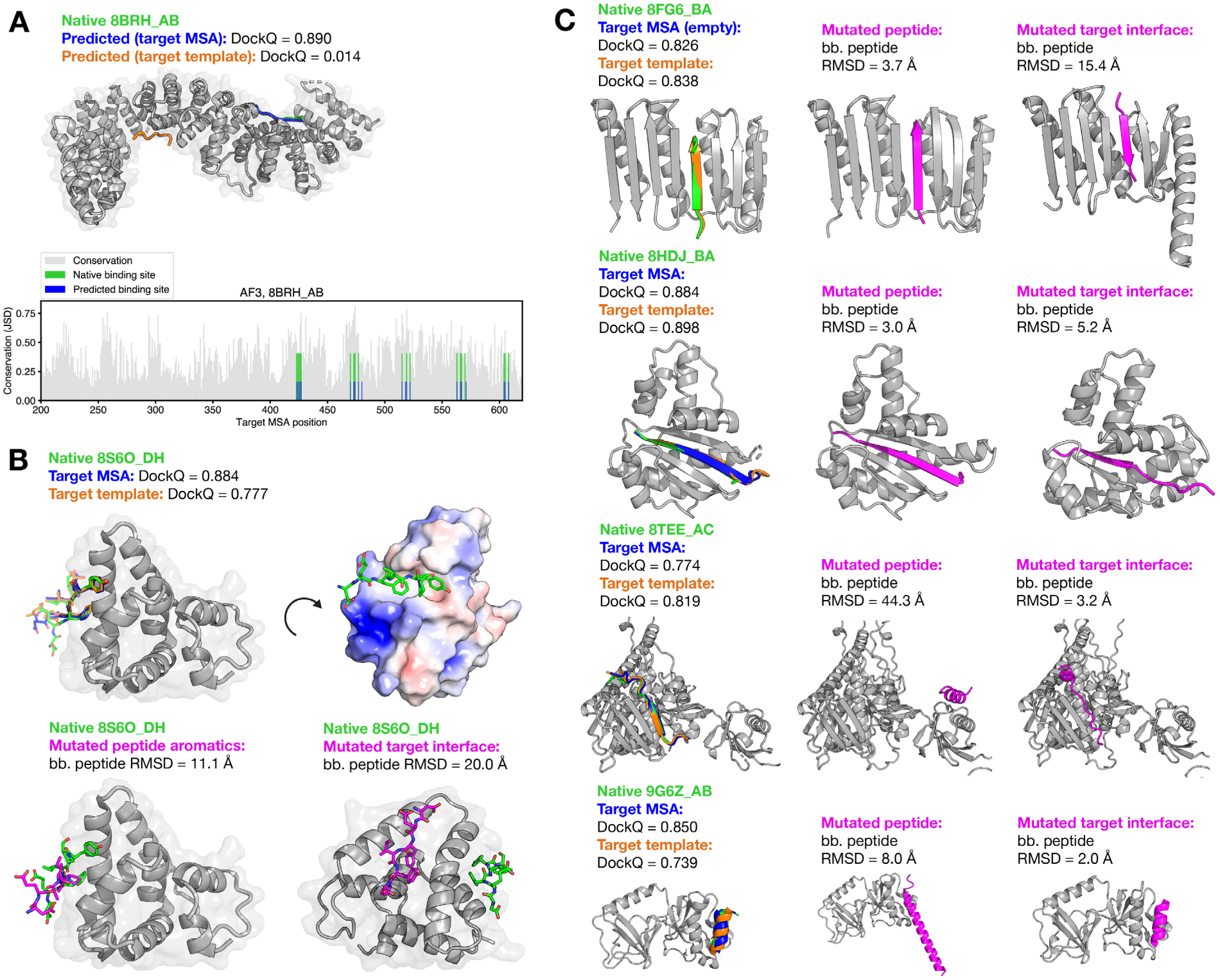
**(a)** top: AF3 predictions for 8BRH_AB made with target MSA input and target template input. Only the native target structure is shown for clarity. bottom: Positional conservation of target residues with native and predicted binding sites overlaid. Only 8BRH_A positions 200-620 are shown for clarity. The prediction shown is with target MSA input. **(b)** top left: AF3 predictions for 8S6O_DH made with only target MSA input and only target template input; top right: A rotated view of the native structure with the electrostatics shown on the target surface; bottom left: AF3 prediction for 8S6O_DH where aromatics in the peptide were mutated to alanine; bottom right: A rotated view of the AF3 prediction for 8S6O_DH where target residues corresponding to the native interface were mutated to alanine. **(c)** AF3 predictions for 8FG6_BA, 8HDJ_BA, 8TEE_AC, and 9G6Z_AB where “mutated peptide” corresponds to a peptide where interface residues were mutated to alanine and “mutated target interface” corresponds to a target where target interface residues were mutated to alanine. For 9G6Z_AB, only the region corresponding to the solved peptide residues in the native PDB structures are shown except for the mutated peptide prediction, where the peptide corresponding to the whole input peptide sequence is shown.

**Figure S17.**
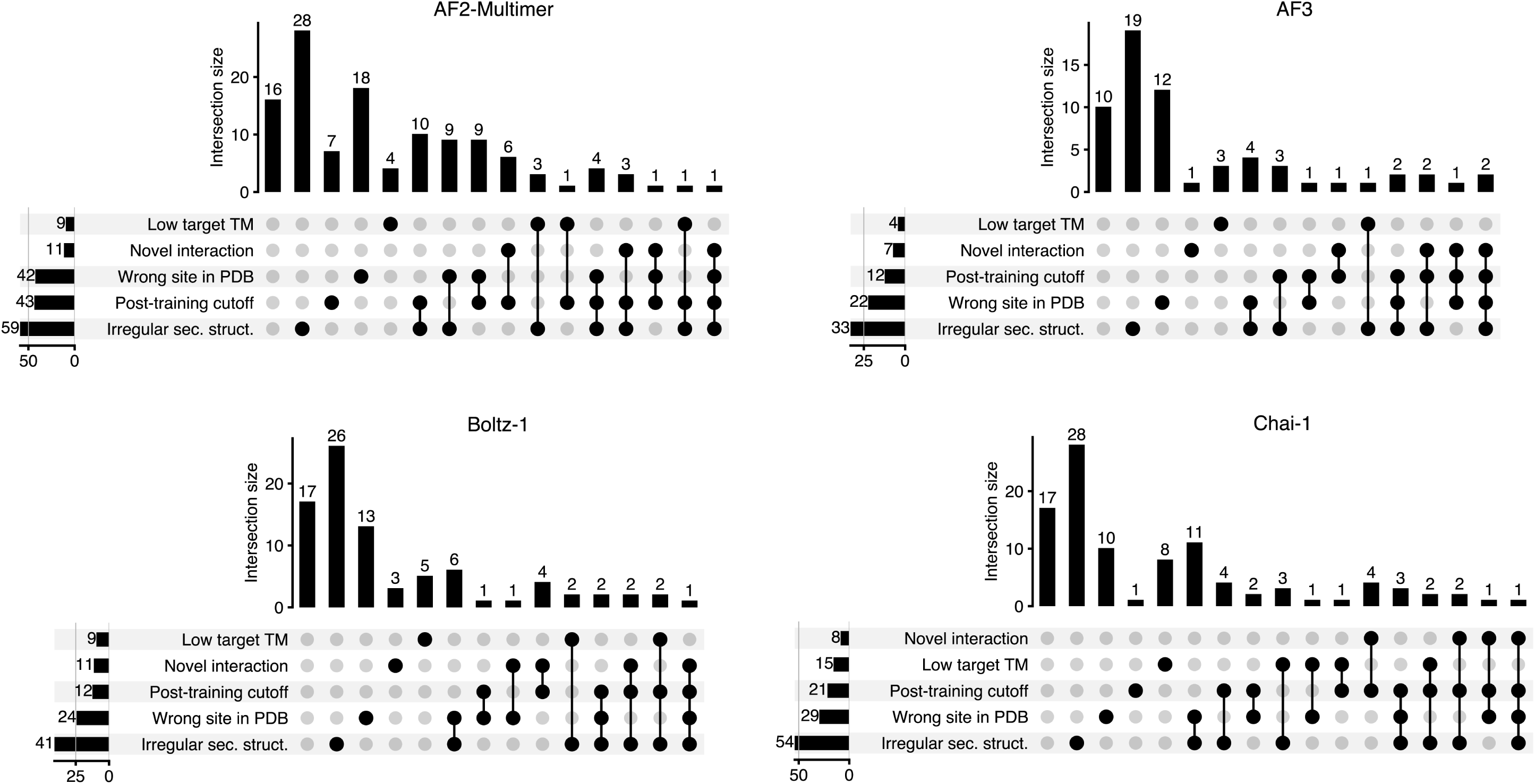
Upset plots for all poor predictions (DockQ < 0.23) under default conditions (unpaired MSA). A TM score *≤* 0.6 is considered poor. A novel interaction is one with 0 training set matches. Structures released past the post-training cutoff are based on the training date cutoffs of each model. An incorrect prediction with at least one match in the training set has “Wrong site in PDB.” Irregular secondary structure is defined by native interface residues having >50% loop-like residues.

**Table S1.** Comparison of settings and success rates between different studies that benchmarked structure prediction models on protein-peptide complexes.

|  | AF Model | Dataset size | Glycine linker | Separate chains | Delimited peptide | Success definition | Success rate |
| --- | --- | --- | --- | --- | --- | --- | --- |
| Tsaban et al. (2022) | Monomer v2.0 | 96 (Large non-redundant set) | Yes | No | Yes | Interface RMSD < 2.5 Å | 38.3% |
| Tsaban et al. (2022) | Monomer v2.0 | 96 (Large non-redundant set) | No | Yes | Yes | Interface RMSD < 2.5 Å | 51% |
| Ko and Lee (2021) | Monomer v2.0 | 203 (PepBDB) | Yes | No | Yes | Peptide RMSD < 2.0 Å | 51% (best of 5) |
| Ko and Lee (2021) | Monomer v2.0 | 203 (PepBDB) | Yes | No | Yes | Peptide RMSD < 2.0 Å | 44% (highest confidence) |
| Bret et al. (2024) | Multimer v.2.2 | 42 | No | Yes | No | DockQ > 0.23 | 40% |
| Bret et al. (2024) | Multimer v.2.2 | 42 | No | Yes | Yes | DockQ > 0.23 | 76% |
| Johansson-Åkhe and Wallner (2022) | Multimer v.2.2 | 112 | No | Yes | Yes | DockQ > 0.23 | 59% |
| Lee et al. (2024) | Multimer v2.2 | 136 | No | Yes | Yes | DockQ > 0.23 | 88% |
| Zhou et al. (2025) | Multimer v2 | Unknown | No | Yes | Yes | Fnat ≥ 0.8 | 53% |
| Zhou et al. (2025) | AlphaFold3 | Unknown | No | Yes | Yes | Fnat ≥ 0.8 | 76.8% |
| Zhou et al. (2025) | Protenix | Unknown | No | Yes | Yes | Fnat ≥ 0.8 | 80.8% |
| Zhou et al. (2025) | Boltz-1 | Unknown | No | Yes | Yes | Fnat ≥ 0.8 | 71.7% |
| Zhou et al. (2025) | Chai-1 | Unknown | No | Yes | Yes | Fnat ≥ 0.8 | 78.8% |

## Methods

### Dataset

We queried the PDB on December 12th, 2024, for structures composed of *≥* 2 chains in the biological assembly, no DNA- or RNA-containing structures, and structures solved with resolution below 2.5 Å. Further filtering was done using a combination of Biopython and PyMol Python API, with filtering steps largely inspired by the curation protocol used to create the Propedia database [13, 14, 15, 16]:

1. The peptide chain must have *≥* 5 and *≤* 50 residues and the target chain must have *≥* 60 residues.
2. At least one peptide atom must be at a distance of 6Å from any receptor atom.
3. The protein–peptide complex must have an interface area of >100 Å^2^.
4. There are no cofactors or ligands within 6 Å of peptide. Common solvents and salts were not considered cofactors or ligands.
5. The peptide has one or fewer non-canonical amino acids.
6. For filtering out interactions mediated by a third protein:

a. There is no other chain making a >100 Å^2^ interface with the peptide.
b. There is no other chain making a >500 Å^2^ interface with the protein.
7. There are no crystal contacts engaging in a >500 Å^2^ interface with the peptide.

To remove duplicates, cluster representatives are used with clusters generated with mmseqs2 at 95% query (protein+peptide) sequence identity [17, 18].

### Base predictions

For benchmarking, we used the LocalColabFold [19, 20] implementation of AF2-Multimer v2.2, AlphaFold3 v3.0.1, Boltz-1 commit d99ceaa61f3af29a4feabee063e5bba8b3f95eb8, and Chai-1 commit 71eff6ac945db726e1d18f4c9e70fba85cd49688. AF2-Multimer v2.3 was not used to enable comparison to other studies benchmarking AF2-Multimer on protein-peptide complexes. Chai-1 was used without ESM embeddings to enable comparison to the other models in the absence of an MSA. An MSA for each chain was constructed with the mmseqs2 server. These MSAs were reused for prediction with Boltz-1 and Chai-1. All models were run with 3 recycling steps, and Boltz-1 and Chai-1 were run with 200 diffusion timesteps (the default setting). AlphaFold3 was run with default settings. DockQ was calculated using DockQ v2.8 All-atom peptide RMS and interface RMS values were calculated with a modified version of the DockQ v2 repo that calculates all-atom DockQ: https://github.com/lindseyguan/DockQ_allatom. DSSP v3.1.4 was used to calculate secondary structure [21, 22]. Hydrophobicity of binding sites was determined by the average Kyte-Doolittle index of hydrophobicity value for binding-site residues [23].

### Inter-chain attention masking

Inter-chain attention masking was implemented in a fork of the OpenFold repository [24]: https://github.com/ lindseyguan/openfold

### Templates

For rigid-body docking, templates were created from the native complex structures. New PDB files were created for each chain, with the entire chain randomly rotated and translated using the Python PyMol API to ensure that models cannot extract the relative docked positions from template coordinates [14].

### Peptide sequence-masked predictions

Masked peptide predictions were made by replacing all peptide residues with the masked token ‘X’ or ‘G,’ keeping the length of the original peptide. The contact maps for AF2-Multimer predictions were extracted from the distograms. Peptide RMSD for Chai-1 and Boltz-1 was calculated from the output structure using the Python PyMol API [14]. RMSD was calculated on only backbone coordinates to enable comparison to the masked peptide residues, which lack sidechains in the model output. The scaling factor was based on parameters used in the DockQ calculation.

### Constrained predictions

For Boltz-1, the binding site was defined as target residues in the native structure containing any atom within 4 Å of any peptide atom. Three binding-site residues were randomly selected to be pocket input. For Chai-1, one target residue-peptide residue pair less than 4 Å apart in the native structure was randomly selected to have a restraint of maximum distance 10 Å.

### Pre-training and post-training cutoff split

For post-training examples, we used all 74 structures from our test set released after the latest training cutoff date, which was 2021-09-30, for AF3 and Boltz-1. The pre-training set consisted of the latest 74 structures from before the earliest training cutoff, which was 2018-04-30, for AF2-Multimer.

### Search for interaction matches in the PDB

For each complex, we first looked for similar instances of the target in the PDB using structure similarity with foldseek [25] and sequence similarity with mmseqs2. The queries were, respectively:

~~~
foldseek easy-search QUERIES_DIR pdb result tmp --exhaustive-search --alignment-type 1
-c 0.8 --cov-mode 2 --tmscore-threshold 0.6
mmseqs easy-search QUERIES.fa pdb_seqres.fa result tmp -c 0.8 --cov-mode 2
~~~

We then checked if there was significant overlap between any binding interfaces in the hit and the binding interface in the query complex structure using TMalign and Biopython [26]. Pythonic pseudo-code for the script used to identify binding site matches from the hits:

~~~
for each complex:
   query = target structure from complex
   matches = []
   for each foldseek/mmseqs hit (’hit’) for this receptor:
        check that this hit was released before the relevant training date cutoff
        query_to_hit_mapping = TMalign(query, hit)
        putative_binding_site = hit residues aligned to query binding site

        binding_site_overlap = 0
        for residue in putative_binding_site:
           nearby_atoms = BiopythonNeighborSearch(residue)
           if nearby_atoms are on another chain:
                # We consider this residue to be part of a binding site
                binding_site_overlap += 1

        binder_rmsd = backbone_RMSD(query_peptide, hit_peptide)

        # Set some cutoffs to indicate a highly similar binding site
        # We use 5 overlapping binding site residues
        # and < 10 Ang binder RMSD
        if binding_site_overlap >= 5 and binder_rmsd < 10:
             matches.append(hit)
~~~

### Mapping peptide sequences to the full-length protein

Peptide chains of each test complex were mapped to a UniProt accession ID using the SIFTS database [27], and bulk UniProt sequences were downloaded using the ID-mapping tool: https://www.uniprot.org/id-mapping. UniProt sequences where the peptide region had > 40% mismatch to the PDB-delimited peptide were excluded. Paired MSAs were generated using original target and the UniProt sequence of the peptide adding either 50 or 100 residues of context around the peptide region. For the prediction, the MSA was sliced to the region corresponding to the PDB-delimited peptide. MSAs were generated using the mmseqs2 server. Predictions were made on the paired MSAs of depth at least 50 and average Jensen-Shannon divergence of the peptide region of at least 0.3.

### Visualization

Upset plots were created with https://github.com/jnothman/UpSetPlot [28]. Protein structure images were created with PyMol v2.5.0 Open-Source.

## Notes

### Competing Interest Statement

The authors have declared no competing interest.

### Funding

- National Institutes of Health: T32GM087237
- National Science Foundation

